# Chromatin Accessibility Plays a Key Role in Selective Targeting of Hox Proteins

**DOI:** 10.1101/473850

**Authors:** Damiano Porcelli, Bettina Fischer, Steven Russell, Robert White

## Abstract

Hox protein transcription factors specify segmental diversity along the anterior-posterior body axis in metazoans. Understanding the basis of Hox function has long faced the problem that, while the different members of the Hox family show clear functional specificity in vivo, they all show very similar binding specificity in vitro. Based on in vitro studies, cofactors may increase Hox binding selectivity however a satisfactory understanding of in vivo Hox target selectivity is still lacking.

We have carried out a systematic analysis of the in vivo genomic binding profiles of all eight *Drosophila* Hox proteins using transient transfection in Kc167 cells to examine Hox protein targeting. We find that Hox proteins show considerable binding selectivity in vivo in the absence of the canonical Hox cofactors Extradenticle and Homothorax. Hox binding selectivity is strongly associated with chromatin accessibility; binding sites in less accessible chromatin show the highest selectivity and the different Hox proteins exhibit different propensities to bind less accessible chromatin. High Hox binding selectivity is also associated with high affinity binding regions, leading to a model where Hox proteins derive binding selectivity through an affinity-based competition with nucleosomes. Provision of the Extradenticle/Homothorax cofactors generally leads to an increase in the number of Hox binding regions and promotes the binding to regions in less accessible chromatin, however the provision of these cofactors has little effect on the overall selectivity of Hox targeting.

These studies indicate that chromatin accessibility plays a key role in Hox selectivity and we propose that relative chromatin accessibility provides a basis for subtle differences in binding specificity and affinity to generate significantly different sets of genomic targets for different Hox proteins. We suggest that this mechanism may also be relevant to other transcription factor families.

## Introduction

Although in vitro studies of transcription factor-DNA interactions have provided extensive insight into how transcription factors bind DNA [1–3], we have less understanding of the basis of transcription factor specificity in the context of chromatin, the environment in which they operate in vivo. Our lack of understanding of in vivo transcription factor specificity is exemplified by the generally poor correspondence between in vivo binding sites identified by Chromatin Immunoprecipitation (ChIP) approaches and predicted target sites based on motifs defined by in vitro studies [4]. Further investigation of the interaction between transcription factors and chromatin is needed to increase our understanding of in vivo transcription factor specificity and improve our ability to predict genomic targets.

A particularly clear example of our insufficient understanding of in vivo targeting of transcription factors is provided by the Hox class of homeodomain proteins. This highly conserved family of transcription factors direct the development of different segmental morphologies along the metazoan anterior-posterior axis, with the classic example of the *Drosophila* Hox gene *Ultrabithorax* (*Ubx*) specifying development of the small rounded haltere balancer organ in the third thoracic segment which, in the absence of *Ubx*, develops as a wing (reviewed in [5–7]). Each of the eight *Drosophila* Hox genes directs the development of a different segmental morphology in vivo. In contrast, all of the Hox proteins show very similar DNA binding preferences when assayed in vitro (reviewed in [8]). A potential way out of this conundrum is provided by the cofactors Extradenticle (Exd) and Homothorax (Hth) in *Drosophila*, and their vertebrate homologues the Pbx and Meis proteins, which interact with Hox proteins to form a tripartite complex [9–12]. In the presence of these cofactors, Hox proteins show a longer consensus binding site and there is evidence of increased differential binding specificity for different Hox proteins [13–15]. In some cases, formation of the Hox-cofactor complex changes the binding preference of the Hox protein providing “latent specificity” [16]. However, we still do not have a satisfactory understanding of in vivo Hox specificity since, first, it is not clear whether the cofactor-enhanced specificity is sufficient to explain the in vivo targeting of Hox proteins and second, in some situations, such as the classic specification of haltere development described above, Hox proteins function in the absence of Exd/Hth cofactors [17].

Previously, we investigated the binding of selected Hox proteins in the context of chromatin through ChIP followed by high-throughput sequencing (ChIP-Seq) in Kc167 cells [18]. We found a strong influence of chromatin state on Hox binding with Ubx and Abdominal-A (Abd-A) binding almost exclusively to DNase1 accessible chromatin, whereas Abdominal-B (Abd-B) exhibited a different specificity and bound to additional genomic sites. This binding, in the absence of Exd/Hth, demonstrated the ability of Hox proteins to exhibit target specificity in the context of chromatin. In addition, the Abd-B-specific binding sites were predominantly in relatively DNase1 inaccessible chromatin. This suggested that histones, rather than simply forming a block to Hox protein binding, restricting the genomic sequence available for binding, might instead play a role in Hox specificity enabling Abd-B to bind to a distinct set of targets through its ability to compete with nucleosomes.

In this report, we present a more comprehensive analysis of the binding of all eight *Drosophila* Hox proteins in the context of chromatin. We demonstrate that they each show distinct chromatin accessibility profiles and that high selectivity of Hox binding is associated with relative inaccessible chromatin. In addition, we find that a major role of Exd/Hth cofactors is to promote Hox binding to relatively inaccessible chromatin. Overall, our studies indicate a key role for chromatin accessibility in determining the selective in vivo targeting of the different members of the Hox protein family.>

## Results

### Hox Protein Binding in Kc167 Cells

We carried out a systematic in vivo analysis of the genome-wide binding of all eight *Drosophila* Hox proteins using our previously established approach [18] designed to maximise comparability between samples. Briefly, we used transient transfection of *Drosophila* Kc167 cells with inducible expression constructs producing Hox-GFP fusion proteins. The cells were fixed 4 h after expression induction and then we used a Fluorescence-Activated Cell Sorter to select cells expressing the same level of Hox-GFP fusion protein. Genome-wide binding profiles were then generated by Chromatin Immunoprecipitation, using an antibody against the GFP tag, followed by high-throughput sequencing (ChIP-Seq). For each Hox protein we collected at least two biological replicates for subsequent analysis.

The binding profiles (Fig 1) show that all eight Hox proteins have distinct but overlapping sets of genomic binding targets. There is a large variation in the numbers of binding regions identified for the different Hox proteins (846 for Antp to 5,685 for Abd-B at q1e-10) and also in the proportion of binding regions unique to an individual Hox protein (Fig 1D). Notably, the centrally-expressed Hox proteins, Antp, Ubx and Abd-A, show very few unique sites.

**Figure 1.**
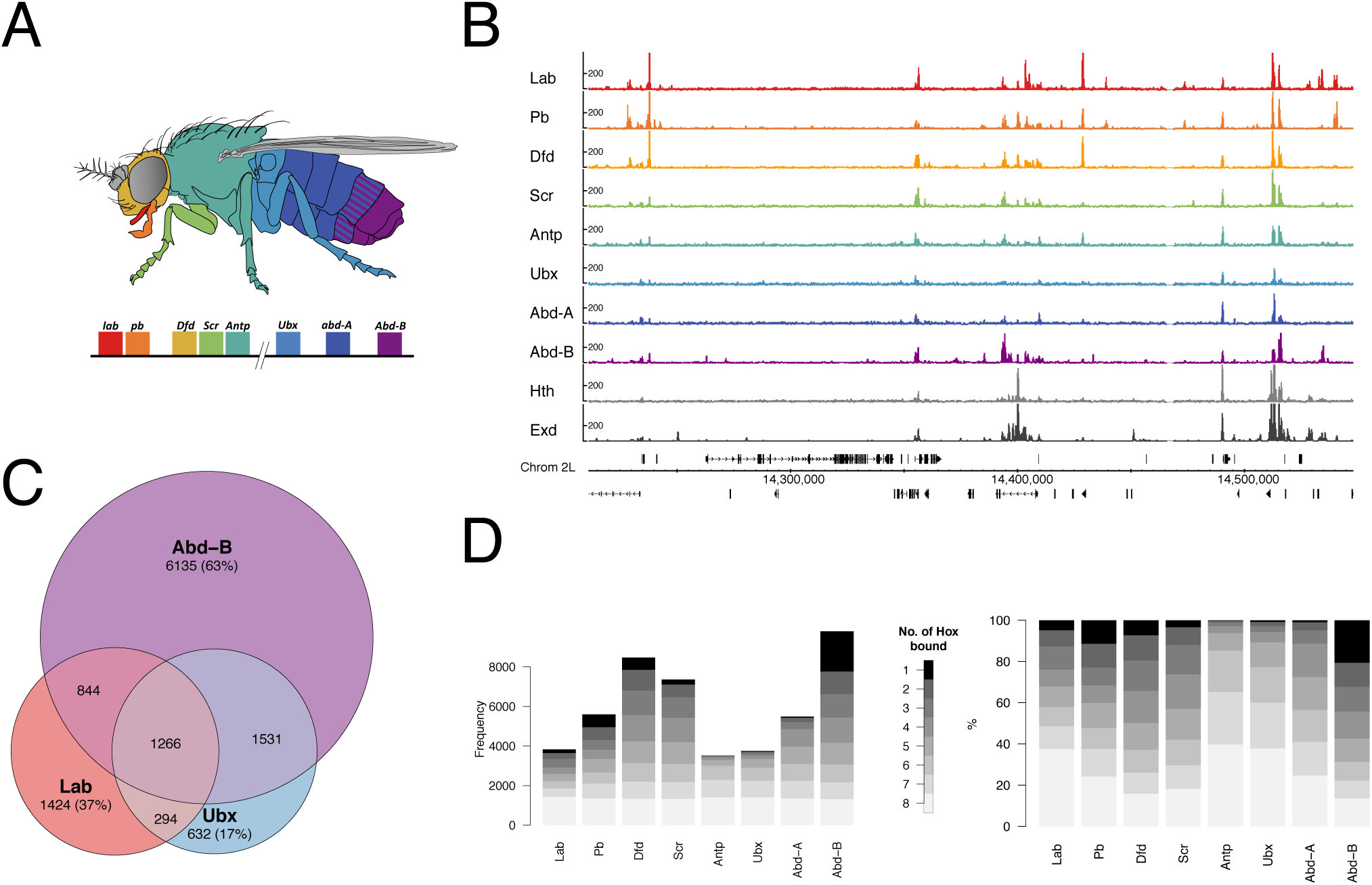
Overview of Hox protein binding in Kc167 cells. **A.** Schematic of an adult Drosophila showing domains of deployment of the 8 Hox genes. **B.** Representative genomic region showing binding profiles of the 8 Hox proteins and the Hox cofactors Hth and Exd. The Exd profile shows the binding of Exd when expressed in association with Hth. **C:** Venn diagram showing overlap analysis of binding regions (q-value 1e-2) for selected Hox proteins Lab, Ubx and Abd-B. Number in brackets gives the number of non-overlapping regions as a percentage of the total number of regions for each protein. **D.** Plots of Hox binding selectivity. For each Hox protein, binding regions (q-value 1e-2) are classified according to the number of Hox proteins bound (see scale). Plotted on the left as frequencies and on the right as percentages, normalised for each Hox protein.

### Motif Enrichment

To investigate the basis for the distinct Hox binding profiles, we compared the enrichment of in vitro-defined Hox binding motifs for the individual Hox proteins in each set of binding regions (Fig 2A). This analysis revealed two insights; first, there is wide variation in the general level of Hox motif enrichment, with an anterior group of Hox proteins, Lab, Pb, Dfd and Scr, showing high enrichments, a central group of Hox proteins, Antp, Ubx and Abd-A, showing little motif enrichment and the most posterior Hox protein, Abd-B, showing substantial enrichment. Second, with the exception of the Abd-B motif, there is little clear discrimination between the different motifs, i.e. for each binding site set the motifs for Lab to Abd-A exhibit rather similar levels of enrichment, whereas the Abd-B motif is discriminating, with low enrichment for the anterior Hox binding sets but high enrichment in the Abd-B binding set. The difference between the Lab to Abd-A motifs and the Abd-B motif fits with a clear shift in base preference in the core motif, from TAAT to TTAT [3,8]. Grouping the motifs on this basis, with Lab to Abd-A motifs grouped as HoxA* and Abd-B as HoxB (Fig 2B,C) provides a simpler view of the enrichment data emphasizing that the three most anterior Hox proteins exhibit much stronger enrichment than the others and demonstrating a clear switch in preference between Lab, the most anterior Hox, and the most posterior, Abd-B. In addition, there is a trend of shift in the HoxA* versus HoxB preference across the whole Hox set.

**Figure 2.**
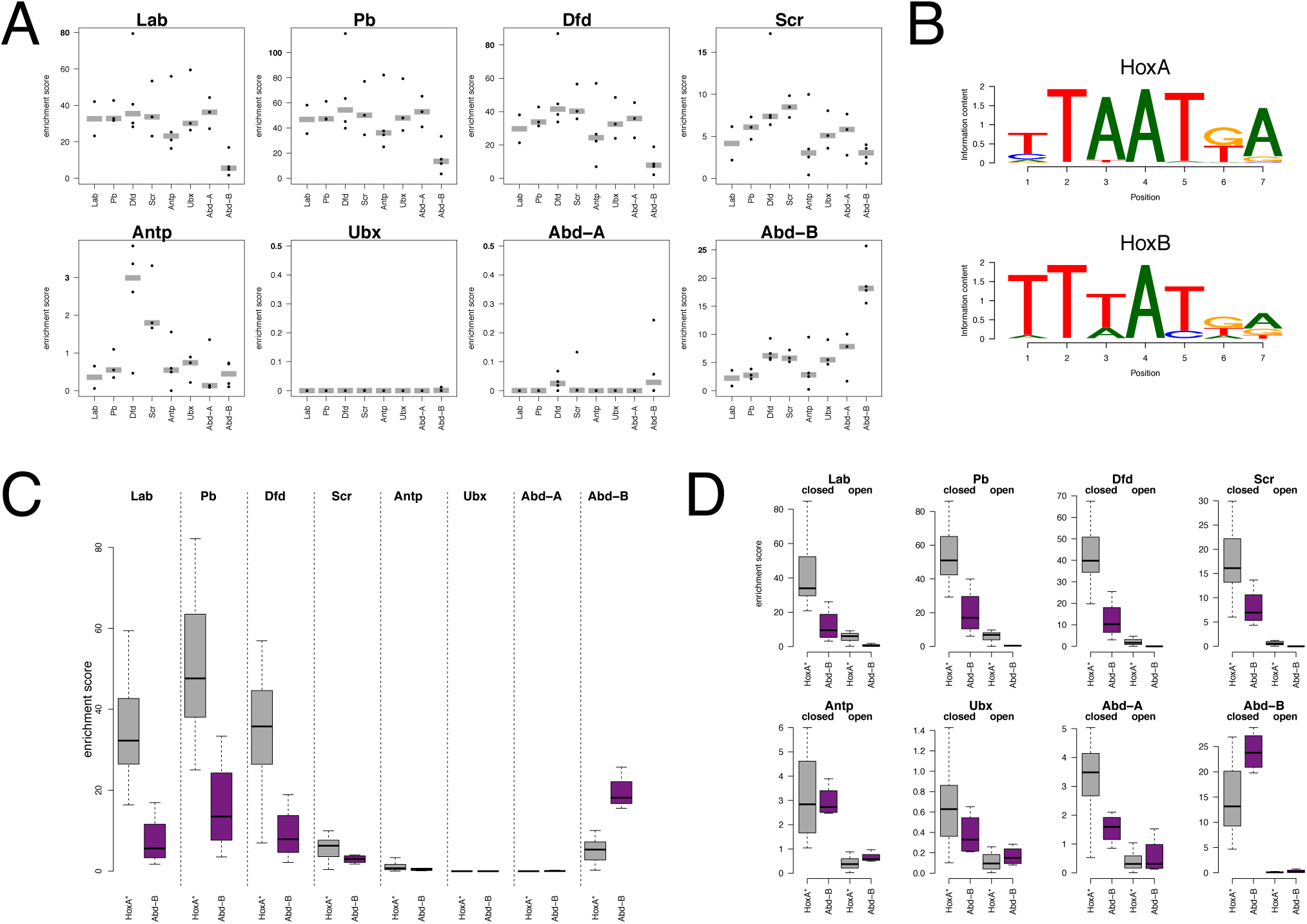
Analysis of Hox binding motifs. **A.** Motif enrichment analysis on the top 500 binding regions (binding summits extended +/- 100bp) for each Hox protein. Plot titles indicate binding region set used and motifs are indicated on the x-axis. Enrichment analysis was performed using PWMEnrich for the Hox motifs in the MotifDb database. Enrichment scores [log_10_(1/p-value)] for individual motifs are indicated (dots) together with the median for each motif set (grey bar). Note the differences in Y-axis scales. **B.** HoxA (consensus motif combining Lab, Pb, Dfd, Scr, Antp, Ubx, Abd-A motifs) and HoxB motifs. **C.** Motif enrichment analysis on the same binding region sets as in (A) using the merged HoxA* (combining scores for the Lab, Pb, Dfd, Scr, Antp, Ubx, Abd-A motifs; grey) and Abd-B (purple) motifs. **D.** Motif enrichment analysis for Hox group peaks - transient separated according to chromatin accessibility, using 500 randomly selected "open" or "closed" regions.

Since we have previously shown that Hox motif enrichment and chromatin accessibility are linked [18], we analysed the motif enrichments separately for “open” and less accessible “closed” chromatin to address the wide variation in general Hox motif enrichment we observe. For this we classified Hox binding regions based on ATAC-Seq scores from untransfected Kc167 cells. We find a dramatic difference in enrichment scores between open and closed chromatin (Fig 2D). Hox binding sites in open chromatin show little enrichment for Hox motifs, whereas high levels of enrichment are found in closed chromatin, particularly for the anterior Hox proteins, Lab, Pb, Dfd and Scr, and for the most posterior Hox, Abd-B. This suggests that the variation in general Hox motif enrichment for the different Hox binding site sets may be linked to the propensity for each Hox protein to bind less accessible chromatin. Accordingly, we examined the chromatin accessibility distribution of the binding sites of the different Hox proteins (Fig 3A) and find a strong concordance with the motif enrichment levels. The Hox proteins with the higher motif enrichments, Lab, Pb, Dfd, Scr and Abd-B bind predominantly to “closed” chromatin whereas those with low motif enrichment, Antp, Ubx and Abd-A, bind predominantly to open chromatin. In addition, the chromatin accessibility distributions show interesting progressions. Anteriorly from Antp and posteriorly from Ubx the Hox proteins present a sequence of increasing binding to less accessible chromatin. These progressions provide an intriguing link between the domains of action of Hox proteins along the body axis and their binding to chromatin.

**Figure 3.**
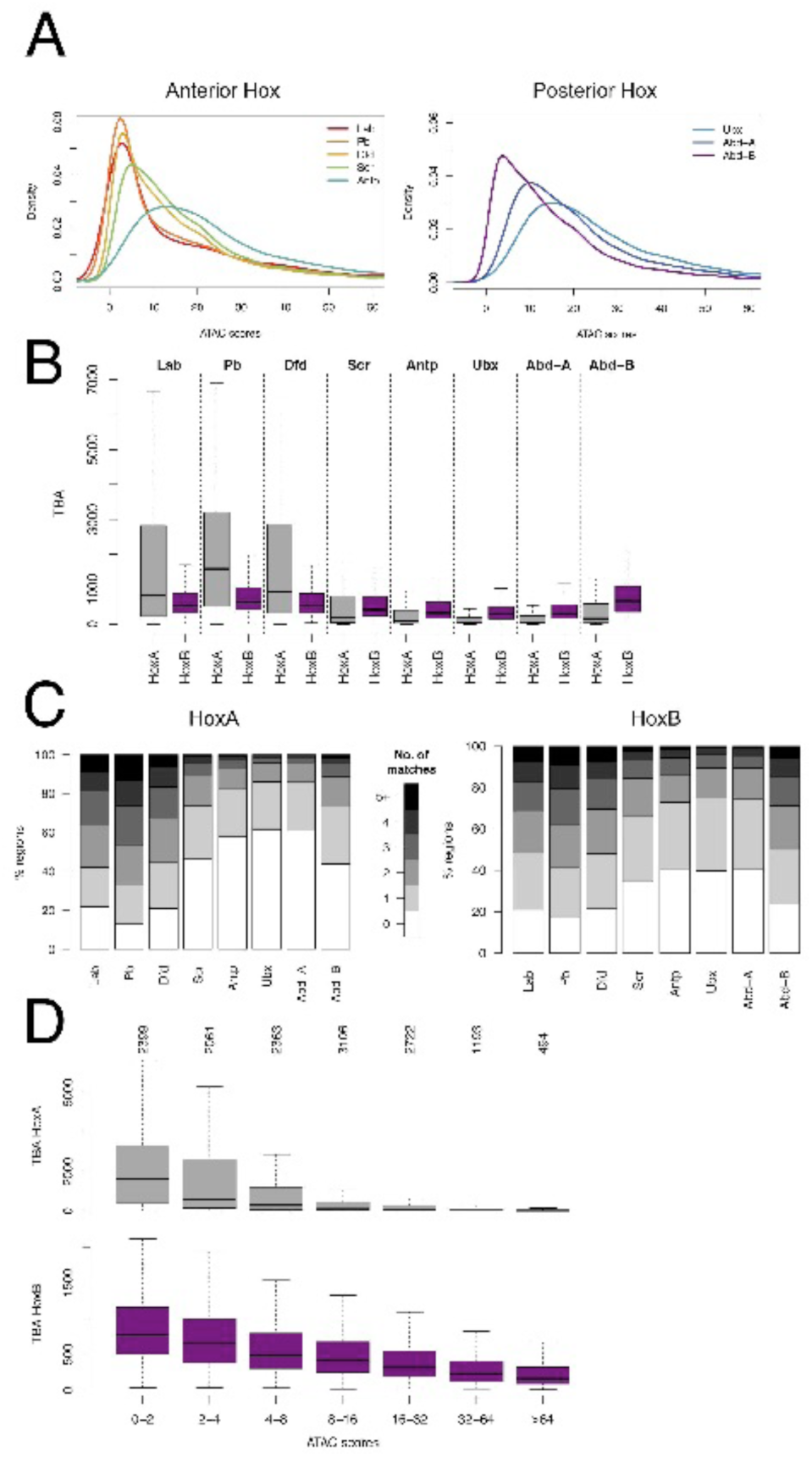
Roles of chromatin accessibility and affinity in Hox binding. **A.** Density plots of mean ATAC-seq scores for Hox group peak regions. Left plot anterior Hox proteins, right plot posterior Hox proteins. **B.** Boxplot of the Total Binding Affinity (TBA) for HoxA (grey) and HoxB (purple) motifs for the top 500 binding regions for each Hox protein. **C.** Plots of Hox group peak regions for each Hox protein classified according to the number of motif matches (HoxA on left, HoxB on right) they contain (see scale). For each Hox protein, matches were counted in the top 500 (by ChlP score) Hox group peak regions using the matchPWM function in Biostrings R package with min.score=90%. **D.** Boxplot showing relationship of TBA and chromatin accessibility, with HoxA (upper) and HoxB (lower) TBA for regions bound by any Hox binned by ATAC score.

Another way to characterize binding sites is through their Total Binding Affinity (TBA; [19,20]), scanning each binding region to produce a cumulative score based on the quality of motif match and the number of motif matches. Similar to the situation with motif enrichment, we see a clear correspondence between TBA and chromatin accessibility distribution. The binding sites for Hox proteins that bind to less accessible chromatin, Lab, Pb, Dfd and Abd-B, show high TBA for their preferred motifs, whereas the binding sites for Hox proteins that bind predominantly open chromatin show low TBA (Fig 3B). The high TBA scores are based on both the quality and number of motif matches in each 200bp binding region (Fig 3C). In addition, the TBA shows a clear switch in Hox motif preference (HoxA versus HoxB) between anterior Hox proteins and Abd-B, as well as a trend in preference switching across the whole Hox set (Fig 3B). The general relationship between TBA and chromatin accessibility is shown in Fig 3D.

Overall, this analysis shows the relevance of both specific binding affinity, based on the quality and quantity of preferred motifs in binding regions, and chromatin accessibility for Hox protein target site selection. Binding to closed chromatin is associated with high TBA that may enable Hox proteins to effectively compete with nucleosomes. In addition, Hox binding occurs across a range of chromatin accessibility and here competition with chromatin may provide the potential for subtle differences in motif preference to generate different target sets for particular Hox proteins.

### Hox Selectivity

We next examined the relationship between chromatin accessibility and the selectivity of Hox binding, as measured by the number of different Hox proteins binding to any particular region. We find a clear relationship, supporting a key role for chromatin accessibility in Hox selectivity. As shown in Fig 4A-C, increasing selectivity is associated with decreasing chromatin accessibility. Sites showing highest Hox selectivity, binding only one member of the Hox protein family, are concentrated in less accessible chromatin whereas sites in open chromatin tend to be poorly discriminating, binding several different Hox proteins. The relationship is gradual with progressive increase in Hox selectivity associated with decreasing chromatin accessibility across the range of ATAC-Seq scores.

**Figure 4.**
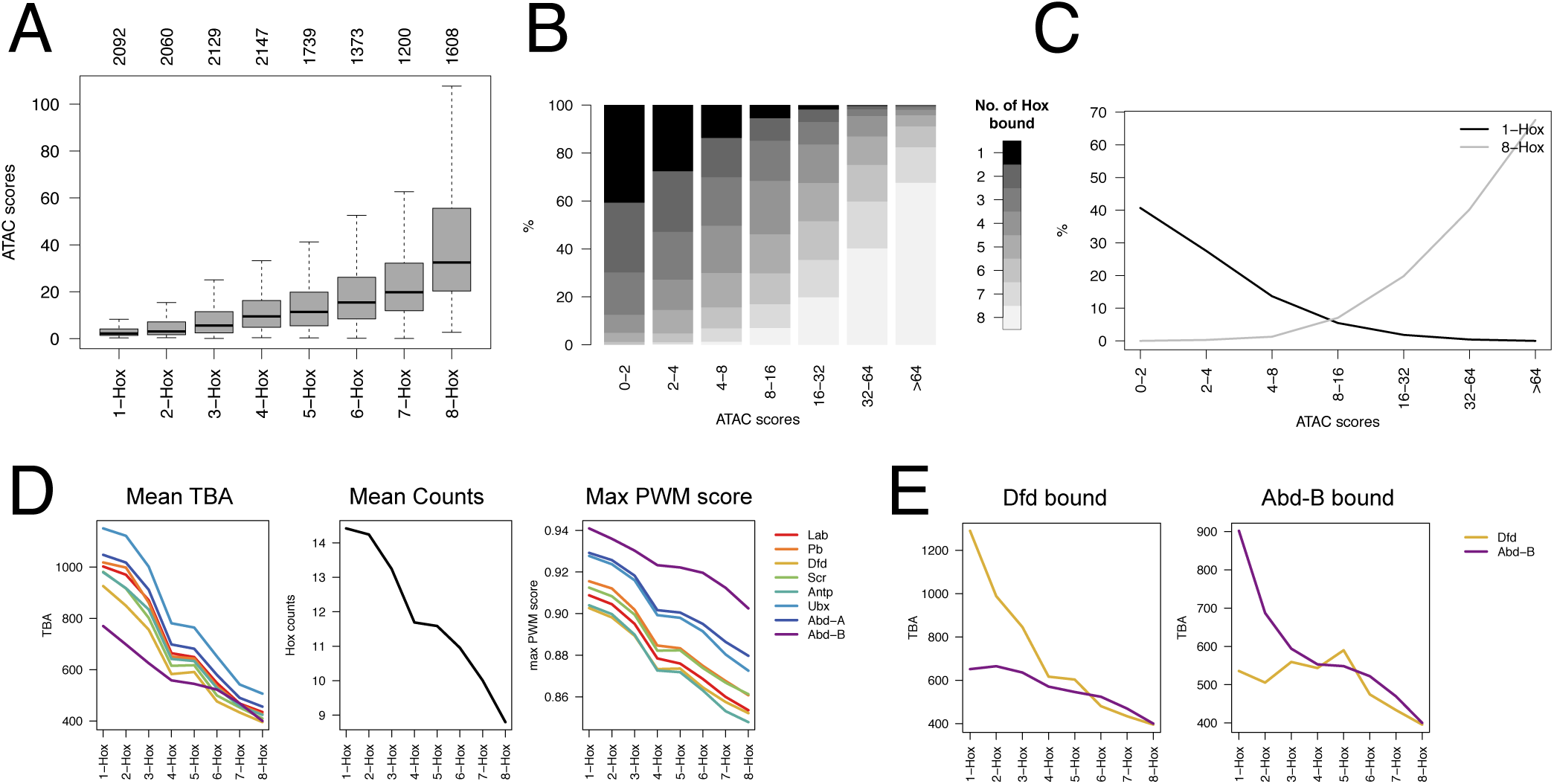
Roles of chromatin accessibility and affinity in Hox selectivity. **A.** Boxplot showing relationship of Hox selectivity to chromatin accessibility (ATAC score). Hox selectivity is represented by binning Hox group peak regions according to the number of Hox proteins bound; 1-Hox = only 1 Hox protein bound, 8-Hox = all 8 Hox proteins are bound. Number of regions in each Hox selectivity bin is shown above the plot. **B.** Plot showing relationship of Hox selectivity classes to chromatin accessibility. The Hox group peaks are then separated into ATAC score bins and the frequency of Hox selectivity classes (see scale) is plotted. **C.** Plot showing opposing distributions of sites binding all Hox proteins (8-Hox) and uniquely bound sites (1-Hox) with respect to chromatin accessibility (ATAC score bins). **D.** Plots showing relationship between Hox selectivity and (from left) TBA (for each jaspar 7-mer Hox PWM), mean number of occurrences of Hox motifs (using matchPWM function with min.score=80%) and highest PWM match score within each region for binding regions as in (A). **E.** Relationship between Hox selectivity and TBA for sets of binding regions for particular Hox proteins. Left for the regions bound by Dfd and right for the regions bound by Abd-B, plotting the TBAs for the Dfd (orange) and Abd-B (purple) motifs.

Hox selectivity is positively correlated with Hox TBA. We see this in the general relationship between Hox selectivity and TBA for the individual Hox binding motifs (Fig 4D). The increasing binding region affinity with increasing Hox selectivity reflects both increasing affinity of individual binding sites (measured by quality of match to Position Weight Matrix (PWM)) and increasing number of Hox binding sites within the binding region. Also, high selectivity for a particular Hox protein is associated with differential TBA for preferred binding motifs. This is illustrated by comparing the target sets for Dfd and Abd-B (Fig 4E). For the low selectivity sites the TBA plots are similar, however at the higher selectivity sites, on the left of the plots, the TBA values show specific inflexion; for Dfd sites there is a specific rise in TBA for the Dfd motif, whereas TBA for the Abd-B motif remains relatively flat. For the Abd-B target set, the reverse occurs. Fig S1 shows a further analysis of the relationship between Hox selectivity and binding region affinity, with subsetting according to chromatin accessibility.

Overall, the association of high Hox selectivity with relatively inaccessible chromatin and high affinity binding regions indicates an interplay between affinity and chromatin accessibility in enabling Hox proteins to bind to different target sets. In a model of competition between nucleosomes and Hox proteins, binding to less accessible chromatin requires a higher affinity interaction between the Hox protein and the binding region. Accordingly, we find Hox binding regions in open chromatin show little discrimination, binding several or all Hox proteins. However, in less accessible chromatin, where competition with nucleosomes provides a basis for selective binding based on affinity, binding regions show more discrimination.

### Roles of the canonical Hox cofactors, Exd and Hth

In many situations, Hox proteins bind in association with the canonical Hox cofactors, Exd and Hth [8,10]. To examine the roles of Exd/Hth in Hox binding we systematically expressed Hth in bicistronic constructs with each GFP-tagged Hox protein and generated ChIP-Seq binding profiles for the Hox proteins as described above. Kc167 cells lack Hth but do express Exd, which is cytoplasmic in the absence of Hth. Expression of Hth recruits Exd into the nucleus and provides Exd/Hth cofactor function [18].

The addition of Exd/Hth generally promotes Hox binding and for all the Hox proteins there is a significant set of cofactor-enhanced regions (Fig 5A). Since we previously showed that Exd/Hth increases the ability of Ubx to bind to closed chromatin [18], we examined the effect of Exd/Hth on the chromatin accessibility distribution. We find that for all Hox proteins, apart from Abd-B, the chromatin accessibility profile is shifted towards lower ATAC-Seq scores indicating that the provision of Exd/Hth enables Hox proteins to bind to less accessible chromatin (Fig 5B,C and Fig S2). Comparing sites where the presence of cofactors results in significantly enhanced binding (cofactor-enhanced sites) with sites that are bound by Hox protein both in the presence and absence of Exd/Hth (common sites) reveals a clear difference in chromatin accessibility. The common sites are predominantly in open chromatin whereas the cofactor-enhanced sites are generally in closed chromatin and for all Hox proteins there is a clear decrease in median ATAC-Seq score for the cofactor-enhanced sites compared with the common sites (Fig 5D).

**Figure 5.**
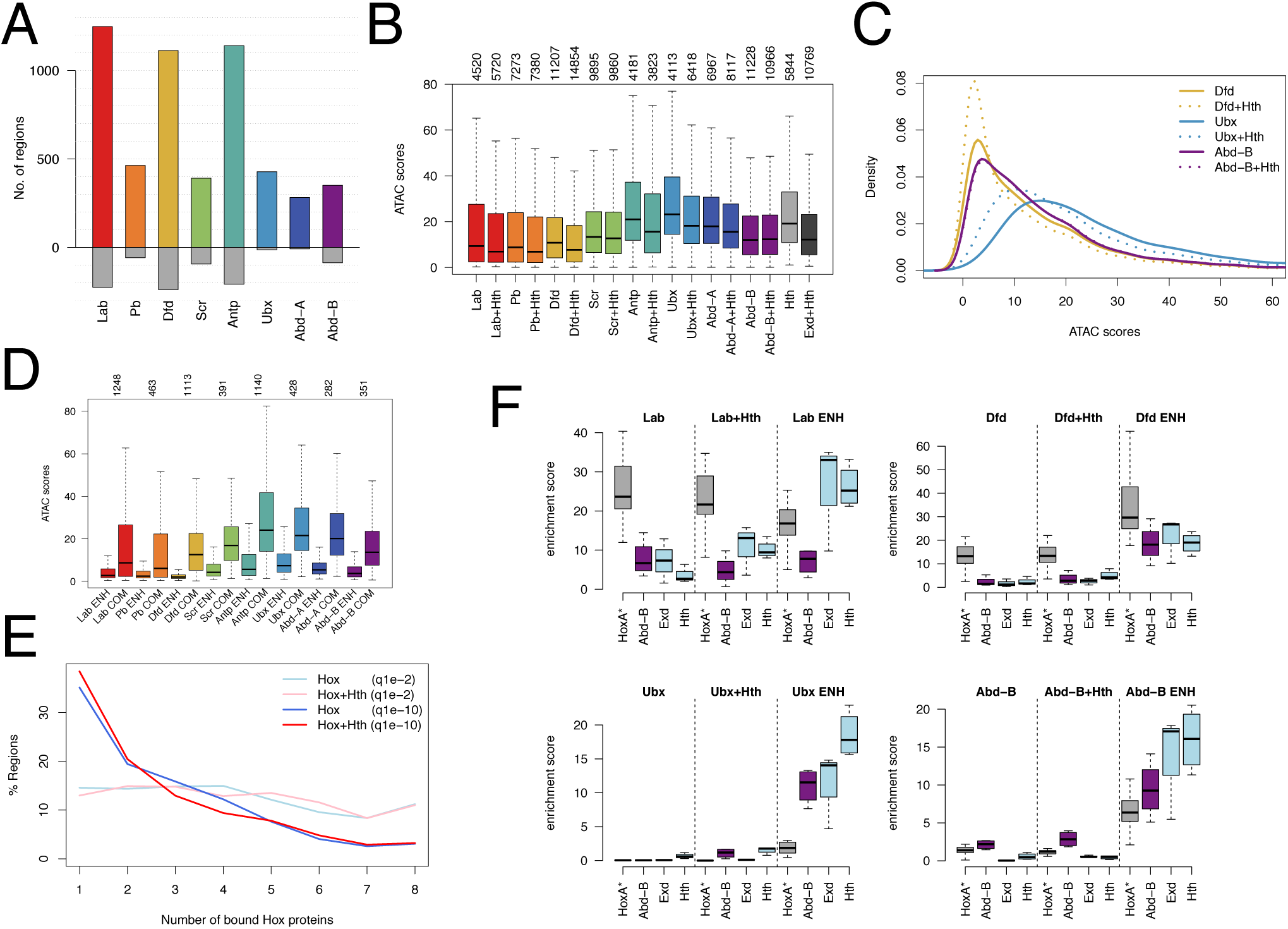
Effect of Exd/Hth on Hox binding. **A.** Plot showing the number of cofactor-enhanced binding regions based on differential binding (fdr <= 0.01, logFC >= 1 and both replicates bound at macs q1e-2). Regions more bound in the presence of Exd/Hth (Hth+Hox) are shown in colour as positive numbers, regions more bound in Hox alone compared to Hth+Hox are shown in grey underneath. **B.** Boxplot of ATAC scores in Hox group peaks for Hox proteins in the absence (Hox) or presence (Hox+Hth) of Exd/Hth. The Hth regions are bound by Hth-GFP and the Exd+Hth regions are bound by Exd-GFP in the presence of Hth. Numbers of bound regions are indicated above the plot. **C.** Density plots of mean ATAC-seq scores for Hox group peaks bound by Dfd, Ubx and Abd-B with and without Exd/Hth showing the effect of the cofactors on the chromatin accessibility profile. Solid lines: Hox alone, dotted lines: Hox in presence of Exd/Hth. **D.** Boxplot comparing chromatin accessibility of Exd/Hth enhanced regions (Hox ENH) versus common regions (bound similarly in the presence or absence of Exd/Hth; Hox COM). Numbers for Hox ENH regions are given above the plot and the same number of randomly selected common regions was used for Hox COM. **E.** Plot showing lack of effect of Exd/Hth on the Hox selectivity profile plotting percentage of regions in each of the Hox selectivity classes for Hox alone (Hox) and in the presence of Exd/Hth (Hox+Hth) for both low and high stringency binding regions. **F.** Motif analysis comparing motif enrichment for binding regions for Hox alone (Hox), Hox in the presence of Exd/Hth (Hox+Hth) and Exd/Hth cofactor enhanced binding regions (Hox ENH) using 500 randomly selected regions from each class for selected Hox proteins Lab, Dfd, Ubx and Abd-B. Motifs are HoxA* (grey; see Fig 2), Abd-B (purple, see Fig 2), and the Exd and Hth motifs (light blue).

Although there is considerable in vitro evidence that Exd/Hth can increase the specificity of Hox binding [13–16], their role in vivo is less clear. As illustrated above, Hox proteins can bind to distinct sets of genomic targets in vivo in the absence of Exd/Hth. Cluster analysis based on ChIP-Seq reads provides a global view, showing that individual Hox binding profiles cluster separately from one another and, strikingly, Hox plus Exd/Hth profiles cluster together with their respective Hox; e.g Dfd and Dfd+Hth cluster together and separately from Lab and Lab+Hth (Fig 6). This demonstrates that Hox proteins display clear individual specificities in vivo independently of Exd/Hth. We also note from this analysis that the anterior Hox proteins Lab, Pb and Dfd show a close association, clustering together and distinct from the remaining Hox proteins which fits with their grouping together on the basis of high motif enrichment in their binding regions (Fig 2C).

**Figure 6.**
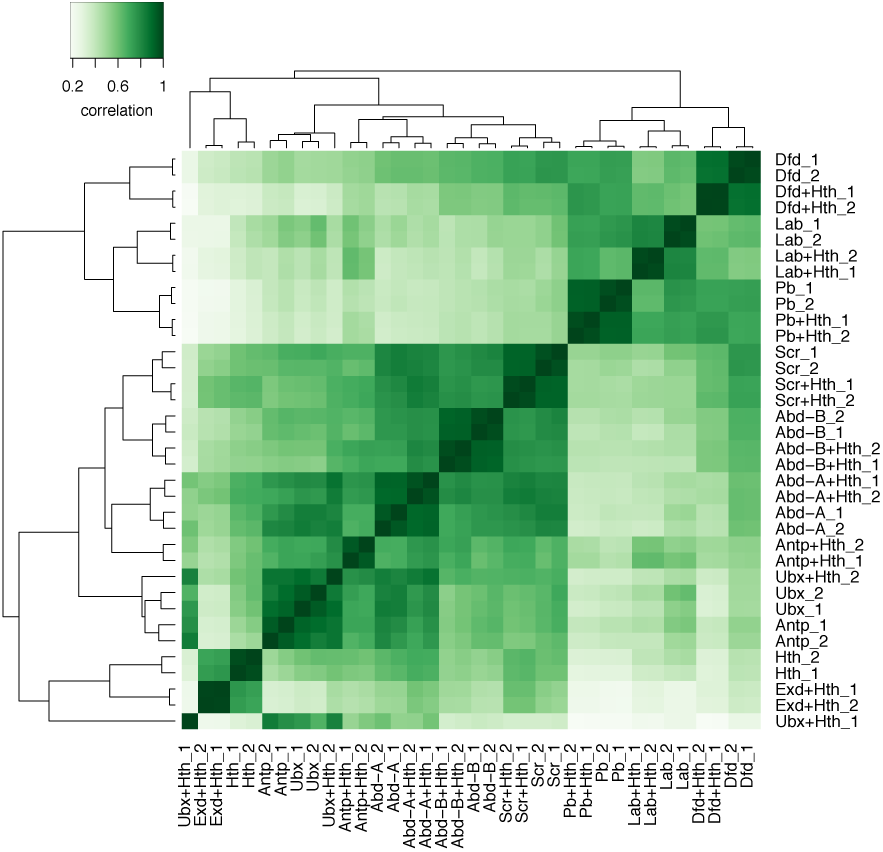
Hox specificity is expressed independently of Exd/Hth. Correlation heatmap of ChlP-Seq reads showing general clustering together of individual Hox and Hox+Hth samples. Reads were counted overlapping 20bp windows of the union of macs q1e-2 bound regions across all Hox and Hox+Hth samples (for details see Materials and Methods).

We assessed the global effect of Exd/Hth on Hox target selectivity by examining the cofactor effect on the distribution of regions according to the number of different Hox proteins they bind. We find that this Hox discrimination profile is little changed by the presence of Exd/Hth (Fig 5E). We also examined the effect of Exd/Hth on the motif enrichment profiles (Fig 5F). In general, the enrichment profile for the Hox motifs is little affected by the addition of Exd/Hox although there are clear increases in the enrichments of Exd and Hth motifs. In the Antp, Ubx and Abd-A target sets the provision of Exd/Hth enhances the relative enrichment of the Abd-B motif above the others and this may represent the latent specificity effect of Hox/Exd dimer binding [16](see also Fig S3).

We performed de novo motif finding analysis on sites where the cofactors significantly increase binding (Fig S4). Combining the most similar motifs, leads to the identification of three classes of cofactor-Hox PWMs (Fig 7A). A k-mer analysis shows that these consensus sequences are the most enriched k-mers in the cofactor-enhanced binding regions (Fig 7B). Preference between these three PWMs provides a clear view of the graded motif preferences across the whole set of eight Hox proteins. Strikingly, these in vivo derived preferences correspond extremely well with the preferences defined by in vitro SELEX analysis of Hox binding in association with Exd [16] (Fig 7C).

**Figure 7.**
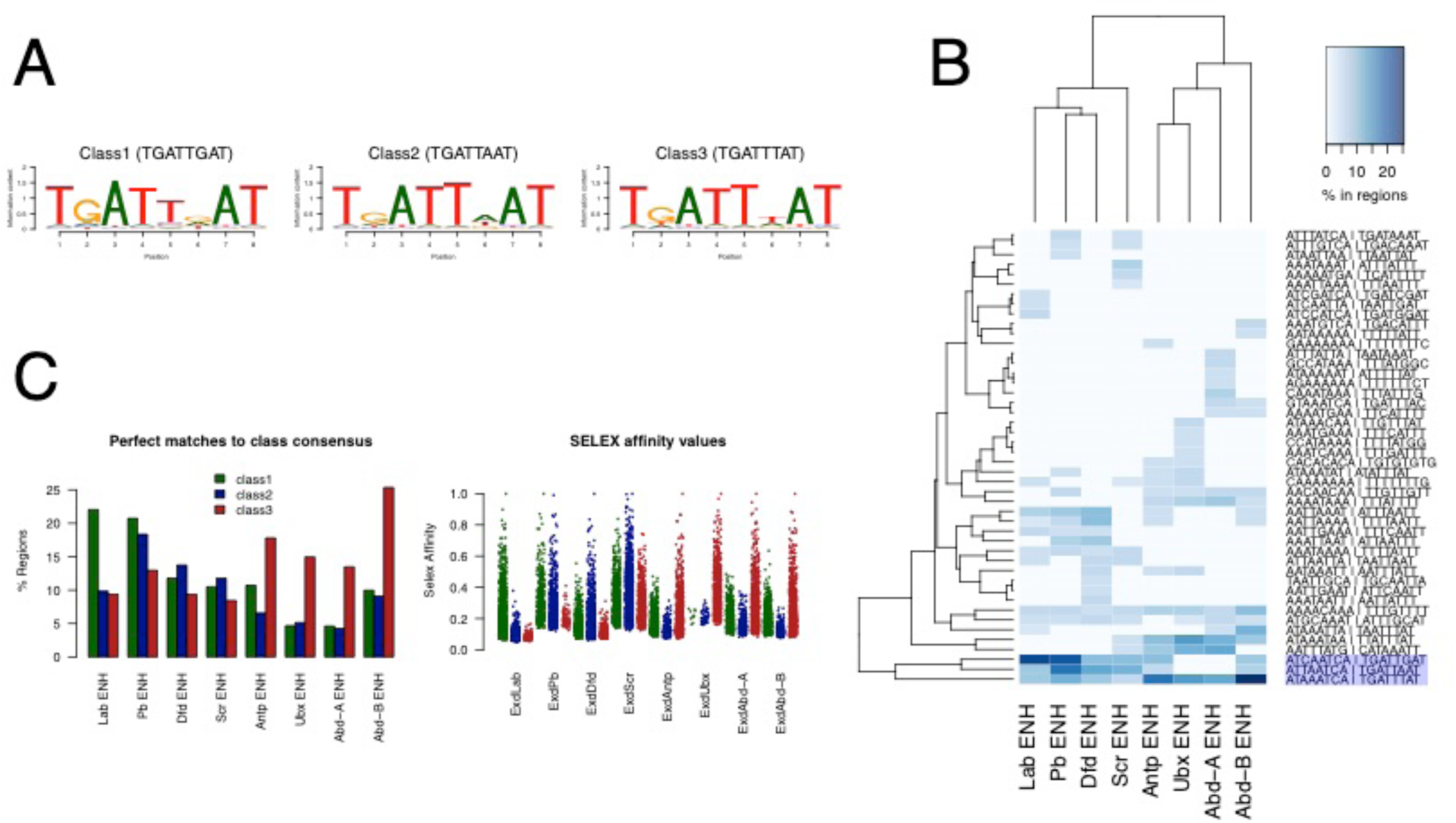
In vivo motif preferences for Hox binding in the presence of Exd/Hth. **A.** Constrained pattern matching on in vivo binding regions defines 3 classes of consensus sequences. Matches to the pattern TGATDAT (where D=A or G or T), based on the in vitro SELEX Exd-Hox sites [16], in defined sets of binding regions were used to create the three matrices. The binding regions used were: Class 1 unique Exd/Hth enhanced Lab bound; Class 2 unique Exd/Hth enhanced Pb or Dfd or Scr bound and Class 3 unique Exd/Hth enhanced Antp or Ubx or Abd-A or Abd-B bound. The pattern matching allowed one mismatch. **B.** The Class 1, 2 and 3 consensus sequences (highlighted) are the most enriched 8-mers in unbiased k-mer enrichment analysis on Exd/Hth-enhanced Hox binding regions (Hox ENH). Enrichment of the top 15 k-mers in each binding region set is plotted as a heatmap. **C.** Correspondence of in vivo and in vitro binding specificities. The left plot shows the percentage of regions from the Exd/Hth-enhanced Hox binding regions with perfect matches to the Class 1, 2 and 3 consensus sequences. The right plot shows the affinity scores for SELEX 16-mers which contain the three Class consensus sequences for the different SELEX Hox+Exd experiments in Slattery et al. [16]; Class 1: green, Class 2: blue; Class 3: red.

Overall, we find that Exd/Hth has significant effects on in vivo Hox binding; for example, almost doubling the number of Dfd-bound regions (increasing from 4782 to 8958 peaks at q1e-10). The cofactors increase the length of the enriched binding motifs and facilitate Hox protein binding to less accessible chromatin.

### Hox binding and chromatin accessibility

To understand better the link between chromatin accessibility and Hox binding, we investigated both the effects of Hox binding on accessibility and also the effects of other transcription factors that either promote Hox binding or are known to be able to open chromatin (so called pioneer factors). To study effects on chromatin accessibility using ATAC-Seq, we generated stable cell lines expressing representative Hox proteins, Dfd, Ubx and Abd-B. We compared the ATAC-Seq profiles of induced versus non-induced cell lines and find clear evidence that Hox proteins vary in their propensity to open chromatin. We see chromatin opening by Dfd and Abd-B but not by Ubx (Fig 8A). Differential peak analysis, confirms that Dfd and Abd-B demonstrate robust chromatin opening with the generation of 430 and 832 significantly enhanced ATAC-Seq peaks respectively (at log fold change >1.5) (Fig 8B and Table S6).

**Figure 8.**
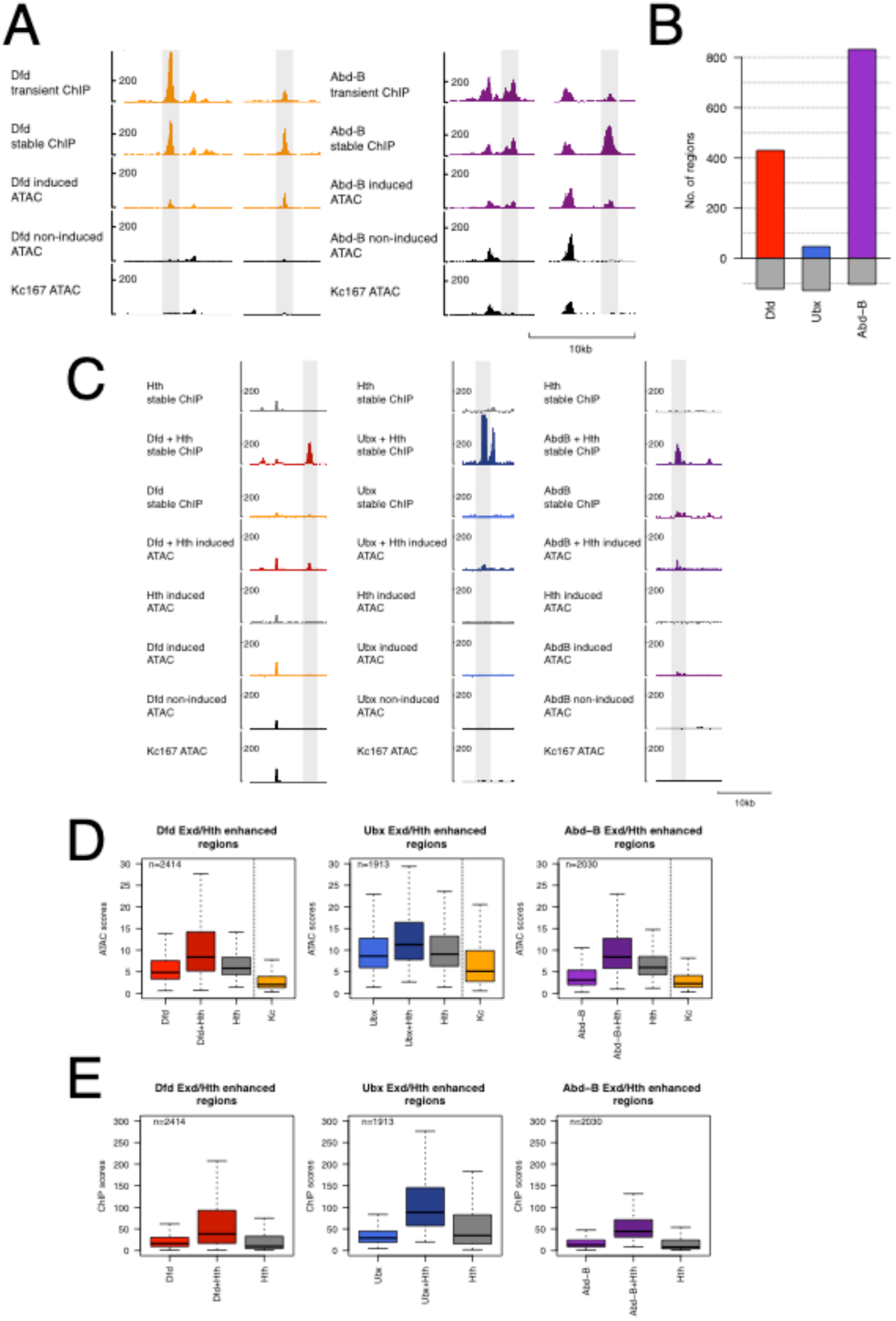
Hox proteins collaborate with Exd/Hth promoting chromatin accessibility. **A.** Representative ChlP-Seq and ATAC-Seq profiles showing increased chromatin accessibility on Dfd and Abd-B binding. **B.** Number of regions with significantly increased chromatin accessibility (edgeR fdr <=0.01 and logFC>= 1.5 for ATAC-Seq reads) on induced versus non-induced samples for Dfd, Ubx and Abd-B are shown as coloured bars. Number of regions with significantly reduced ATAC-Seq reads (edgeR fdr <=0.01 and logFC<= −1.5 for ATAC-Seq reads) are shown in grey as negative values. **C.** Representative ChlP-Seq and ATAC-Seq profiles showing collaboration between Hox and Exd/Hth promoting chromatin accessibility. **D.** Boxplot of ATAC scores in Exd/Hth-enhanced Hox binding regions for stable lines expressing Hox alone, Hox in the presence of Hth (Hox+Hth), Hth alone and, as a reference, the basal Kc167-cell (Kc) ATAC scores. All three Hox+Hth show increased ATAC scores compared to either Hox alone or Hth alone; p-values <0.01, Dunn's Kruskal-Wallis Multiple Comparison. Although the Kc167 ATAC scores cannot be directly compared to the stable cell line ATAC score data, the low median scores indicates that these regions are relatively inaccessible in the basal Kc167 state. **E.** Boxplot of ChlP-seq scores in the same regions as in (D) showing Hox ChlP, Hox Chlp in the presence of Exd/Hth (Hox+Hth) and Hth ChlP.

To investigate the role of Exd/Hth we generated stable cell lines expressing Dfd, Ubx or Abd-B proteins together with Hth and we find that the cofactors promote chromatin opening (Fig 8C). Examining the regions with enhanced Hox binding in the presence of Exd/Hth, we find that addition of the cofactors increases the median ATAC-Seq score in partnership with each of the three Hox proteins (Fig 8D). In contrast, when Hth is expressed in the absence of a Hox protein these regions show low chromatin accessibility and little evidence of Hth binding (Fig 8E), indicating that the Hox proteins and Exd/Hth work in collaboration to open chromatin. Differential peak analysis on the 21,002 Hox group peaks - stable regions, comparing induced versus non-induced for the Hth stable cell line supports the lack of significant chromatin opening when Hth is expressed in the absence of Hox proteins (Table S6).

We directly examined the effect of opening chromatin on Hox binding by co-expressing Hox proteins with the haemocyte lineage-determining factor Glial cells missing (Gcm), which is believed to act as a pioneer factor [21]. Kc167 cells show characteristics of haemocytes, which in the in vivo lineage are induced to differentiate into plasmatocytes by Gcm [22,23]. We first established, by ChIP-Seq and ATAC-Seq in Kc167 cells expressing Gcm-GFP, that Gcm binding is associated with chromatin opening (Fig 9A-C). We then expressed Gcm in conjunction with either Dfd or Ubx and find that the presence of Gcm leads to novel binding sites for both Dfd and Ubx (Fig 9A-B). This provides a direct experimental demonstration of the role of chromatin accessibility in Hox target selection. For Dfd, the provision of Gcm generates 1168 novel sites (at q1e-2, 13% of the total Dfd binding sites in the presence of Gcm), whereas for Ubx, Gcm has a larger effect generating 4291 novel sites (49% of the total Ubx binding sites in the presence of Gcm). The smaller effect of Gcm on Dfd binding may reflect the ability of Dfd to bind to less accessible regions on its own.

**Figure 9.**
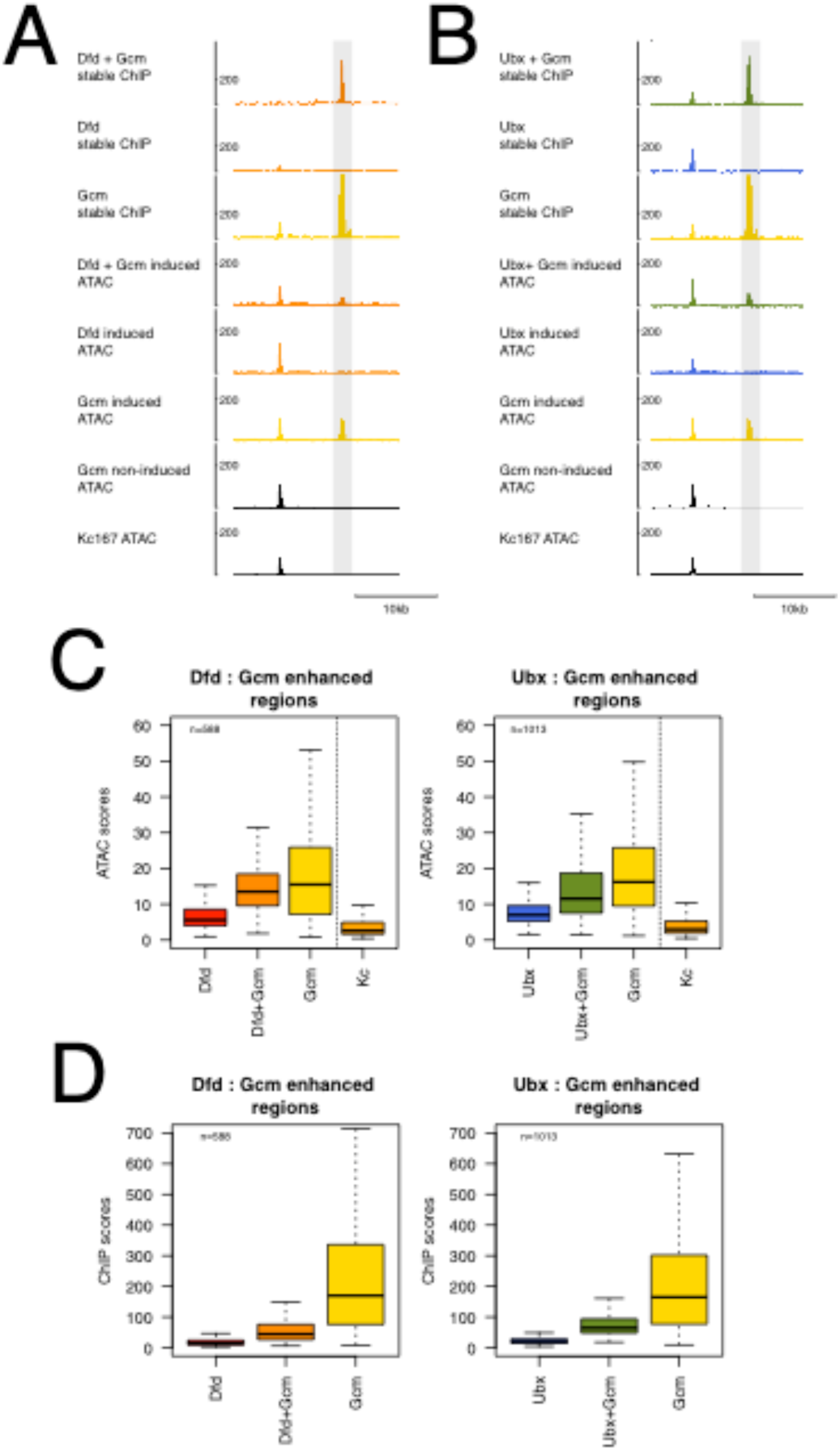
Gcm acts as a pioneer factor promoting Hox binding. **A.** Representative ChlP-Seq and ATAC-Seq profiles showing chromatin opening by Gcm and promotion of Dfd binding. **B.** Representative ChlP-Seq and ATAC-Seq profiles showing chromatin opening by Gcm and promotion of Ubx binding. **C.** Boxplot of ATAC scores in Gcm-enhanced Hox binding regions for Hox, Hox in the presence of Gcm (Hox+Gcm), Gcm and, as a reference, the basal Kc167-cell (Kc) ATAC scores. Both Hox+Gcm show increased ATAC scores compared to Hox alone; p-values <0.01, Dunn's Kruskal-Wallis Multiple Comparison. The high median ATAC scores for Gcm show that these regions are generally open in the presence of Gcm alone. Although the Kc167 ATAC scores cannot be directly compared to the stable cell line ATAC score data, the low median scores indicates that these regions are relatively inaccessible in the basal Kc167 state. **D.** Boxplot of ChlP-seq scores in the same regions as in (C) showing Hox ChlP, Hox Chlp in the presence of Gcm (Hox+Gcm) and Gcm ChlP.

The comparison of the effects of Exd/Hth versus Gcm on Hox binding, reveals two rather different routes to enhance Hox binding. In contrast to the Exd/Hth situation, the sites with significantly increased Hox binding in the presence of Gcm are associated with robust Gcm binding and chromatin opening by Gcm when expressed in the absence of Hox (Fig 9C and Fig S5). Thus, while Exd/Hth and Hox work together to enhance chromatin accessibility, Gcm is able to open chromatin independently of Hox and thereby facilitate Hox binding.

### Comparison of chromatin accessibility and binding site affinity in Hox target selection

To gain an overview of the relationships between accessibility, binding site affinity and Hox occupancy we present the Hox binding data for “open chromatin” as a heatmap of occupancy (percentage of Hox-occupied 200bp open chromatin regions per bin) for regions displayed as a scatter plot of accessibility (logATAC score) versus affinity (logTBA) (Fig 10A, S6). Even within these “open” chromatin regions we see a strong influence of accessibility: regions with low ATAC scores show low occupancy while the most open regions exhibit very high occupancy. In contrast, the correlation between occupancy and TBA is much less strong (Fig 10B). The relevance of relative accessibility for Hox binding is emphasized by the “No Hox” plot where the heatmap illustrates percentage of regions not bound by any Hox protein (Fig 10A). The regions with least accessibility show very high percentages of regions with no Hox proteins bound and followed by a graded decrease in unbound regions as the ATAC scores rise. The strong correlation between Hox binding and chromatin accessibility and the observation that the most open regions show close to 100% occupancy suggests that, while there is a requirement for openness, there is not a requirement for specific binding partners at the bound open regions.

**Figure 10.**
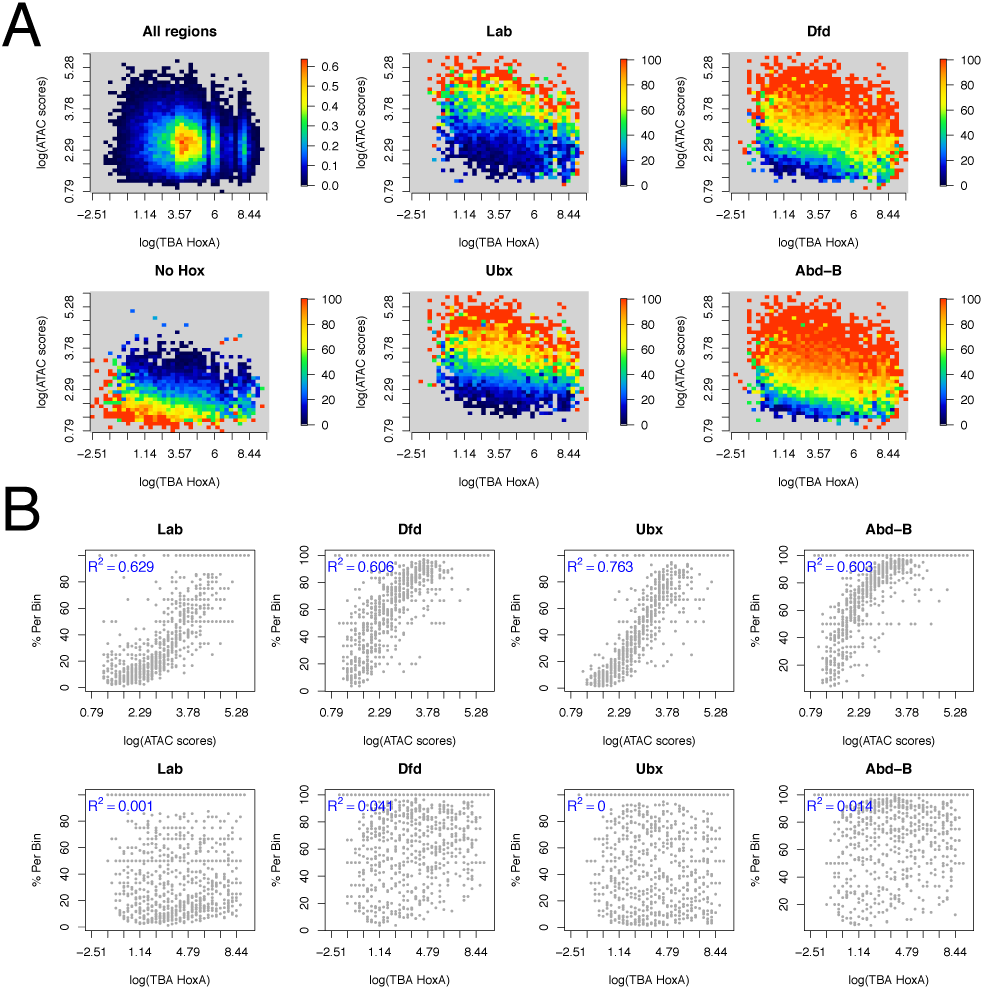
Hox occupancy is more strongly associated with binding region chromatin accessibility than with binding affinity. **A.** Scatter plots of chromatin accessibility (log[ATAC scores]) versus binding affinity (log[TBA HoxA]) for chromatin regions classified as "open". Open chromatin regions were divided into 200bp tiles and the mean ATAC score and TBA for HoxA PWM calculated per tile. The log of these scores was then linearly binned into 40 bins on each axis. For the "All regions" plot the heatmap shows the density distribution. For the other plots, the heatmap shows the percentage of tiles bound by the specified Hox protein per bin or for "No Hox" the percentage of tiles not bound by any Hox protein. The plots are shown for selected Hox proteins and for TBA for the HoxA PWM (for a fuller set of plots including HoxB TBA see Fig S6). **B.** Scatter plots show the strong correlation of occupancy (% per bin) with chromatin accessibility (log[ATAC scores]; upper row) and the poor correlation with binding affinity ((log[TBA HoxA]; lower row). Data as in (A). Further plots in Fig S6.

## Discussion

Ever since the initial analyses of DNA binding by Hox proteins in the 1990s [14,15,24,25], our understanding of the basis for Hox specificity has faced the conundrum that while Hox proteins exhibit clear functional specificity in vivo the different members of the Hox family show very similar DNA binding specificity in vitro. One of the unknowns for our understanding of in vivo Hox specificity has been the effect of chromatin on Hox binding. Our investigations into Hox protein binding in the context of chromatin in Kc167 cultured cells reveal a strong interplay between target selection by Hox proteins and chromatin accessibility. We find that Hox selectivity shows a graded relationship to chromatin accessibility, with sites in relatively closed chromatin showing highly selective Hox binding while sites in open chromatin tend to be unselective and exhibit binding by most or all Hox proteins (Fig 4). The binding regions for different Hox proteins show different chromatin accessibility profiles. Some Hox proteins, such as Dfd and Abd-B bind relatively closed chromatin, whilst others, such as Antp and Ubx, bind almost exclusively to open chromatin (Fig 5). Binding regions in relatively inaccessible chromatin are generally high affinity, with multiple good matches to consensus binding sites, while binding sites in open chromatin tend to have low affinity (Fig 3). Our data fit with a model of in vivo Hox specificity based on the different ability of specific Hox proteins to compete with chromatin and access their DNA binding sites.

The propensity to compete with chromatin could depend on a variety of factors. The link between high TBA and high selectivity indicates that target selectivity could depend on differences in the affinity of interaction with binding sites. Hox proteins with relatively high affinity for their preferred binding sites, and/or with the ability to effectively use multiple binding sites to increase affinity, could compete with chromatin to establish binding at the relatively closed chromatin environments of the selective sites. Hox proteins with high numbers of unique binding regions, e.g. Pb, Dfd and Abd-B would have high affinity for their preferred sites. One the other hand, Hox proteins, such as Antp and Ubx would have lower affinity and be unable to reach the affinity threshold for effective competition with chromatin and so would be restricted to binding less selective open chromatin regions. This could be termed a quantitative affinity model. Alternatively, selectivity could be based on more qualitative differences between Hox proteins, for example they could differ in their ability to bind to nucleosomal DNA, or in their ability to interact with other DNA-binding proteins with whom they could collaborate to compete with chromatin [8,26–28]. A third possibility is that their differential ability to interact with relatively inaccessible chromatin could depend on selective ability to interact with chromatin remodelers to open chromatin at their binding sites.

We have investigated the influence of other DNA-binding proteins on Hox binding in chromatin in two different situations; provision of the canonical Hox cofactors Exd/Hth and provision of the pioneer factor Gcm. The Exd/Hth cofactors physically interact with Hox proteins through binding between Exd and the Hox YPWM motif and other interfaces [29–33]. Provision of Exd/Hth together with individual Hox proteins in Kc167 cells has marked effects on Hox binding, generally resulting in an increase in the number of significant Hox binding sites, promotion of chromatin opening and a shift in the prior chromatin accessibility profile of bound sites towards less accessible chromatin. The Exd/Hth-enhanced Hox binding regions show little Hth binding or chromatin opening when Exd/Hth is expressed in the absence of Hox. Although there is evidence for the vertebrate Exd and Hth homologues acting as pioneer factors at specific sites [34], in our genomic analysis we find little support for pioneer function of Hth but rather Hox and Exd/Hth appear to work together to open chromatin and promote Hox binding. Exd/Hth-enhanced Hox binding regions are strongly enriched in Exd-Hox consensus dimeric binding sites. Overall, the effects of Exd/Hth on Hox binding suggest Exd/Hth provides an increase in binding affinity at the Exd/Hth-enhanced Hox binding sites promoting enhanced competition with chromatin and raising the chromatin accessibility threshold for each Hox protein. The resulting general shift of the chromatin accessibility profile for each Hox protein towards less accessible chromatin fits with the quantitative affinity model. In the second situation, we provide Gcm, a protein that does not physically interact with Hox proteins [26] but which we show has the ability to open chromatin. We tested the effects of the provision of Gcm in conjunction with either Dfd or Ubx. In both cases, the increased chromatin accessibility due to Gcm generated novel Hox binding sites but more for Ubx than Dfd, which fits with the ability of Dfd to bind less accessible regions on its own. In contrast to the situation with Exd/Hth, with Gcm we see no evidence for collaborative effects on chromatin opening. Gcm presents an example of a DNA binding protein that alters the chromatin accessibility landscape thereby affecting Hox binding without necessarily having a direct physical interaction with Hox proteins. This may be a general way that other DNA-binding proteins affect Hox protein targeting through the strong effect of chromatin accessibility on Hox binding, as the almost complete occupancy of the most highly accessible regions (Fig 10) suggests that binding is dependent on accessibility per se without the necessity for interactions with specific partner proteins.

Although Exd/Hth has a strong effect on the number and accessibility of Hox binding regions, we see little general effect of Exd/Hth on the selectivity of Hox binding in vivo. However, particularly for Antp, Ubx and Abd-B the provision of Exd/Hth alters the Hox binding specificity as seen in the increased relative enrichment of the Abd-B motif (HoxB) versus the anterior Hox motifs (HoxA). This may occur through conformational constraints on Hox proteins in the Hox/Exd/Hth complex as in the phenomenon of latent specificity seen in vitro [16].

The specific case of the interaction of Abd-B with Exd/Hth is interesting since Abd-B lacks the YPWM motif, although it may interact with Exd through other interfaces [32], and its binding affinity for DNA in vitro is not increased by Exd [15]. In our data, Abd-B does not follow the same trend as the other Hox proteins in that provision of Exd/Hth does not increase the number of significant peaks detected by Abd-B ChIP. Furthermore, there is no general enhancement in binding to less accessible chromatin, since we observed no decrease in the average ATAC scores for Abd-B binding regions nor any change in the accessibility profile (Fig 5 and S1). These results suggest that Abd-B may not interact with Exd/Hth in vivo. However, further examination of differential binding reveals that there is a significant set of regions where Abd-B binding is enhanced in the presence of Exd/Hth (351 regions at logFC 1 in the transient data set) and in these regions Exd/Hth promotes Abd-B binding to less accessible chromatin (Fig 5A,D). These regions show strong enrichment for Abd-B, Exd and Hth motifs (Fig 5F) and the dimeric Exd-Hox site TGATTTAT is the most enriched motif found by de novo motif analysis on the set of regions that show enhanced binding of Abd-B in the presence of Exd/Hth (Fig S4). Thus Abd-B may interact with Exd/Hth at a subset of sites in vivo. On the other hand, particularly in our stable line data, there is a significant set of regions (1648 regions at logFC 1) that show decreased binding by Abd-B in the presence of Exd/Hth which fits with the antagonism between Abd-B and Exd/Hth described previously [35].

Our data reveal a clear relationship between Hox specificity and binding site affinity (Fig 4). We find that the regions associated with highly selective Hox binding show high TBA, based on multiple binding sites with high scoring matches to Hox consensus binding sites. This fits with the observation that these sites are in less accessible chromatin, suggesting that high affinity is required for effective competition with chromatin. However, this relationship contrasts with the evidence from in vitro SELEX studies and observations at the *ovo/shavenbaby* locus in vivo where highly selective Hox binding is associated with low affinity binding sites [36]. Interestingly, in our studies, while Hox selectivity is linked to high affinity sites over the whole set of binding regions, if we examine the relationship over the subset of regions with higher chromatin accessibility (ATAC scores >25) the relationship is reversed so that higher selectivity is associated with lower TBA (Fig S1C,D). Thus the link between weak binding sites and high Hox selectivity may be applicable in highly open chromatin.

Several features of the binding data for the different Hox proteins, notably the percentage of unique sites (Fig 1D) and the profiles of chromatin accessibility (Fig 3A), show an intriguing graded relationship to the sequence of Hox gene expression along the anterior-posterior body axis. For both these features the central Hox genes represent a minimum state; for example, the ability to access more closed chromatin progressively increases both anteriorly and posteriorly from a low point represented by Antp/Ubx (Fig 3A). These profiles follow the sequence of posterior dominance running posteriorly from Antp to Abd-B [37–39] and are reminiscent of the idea of an evolutionary and developmental ground state represented by the second thoracic segment or the Hox gene Antp [40,41]. This suggests that in building on the ground state the progressive ability of Hox proteins to engage with binding sites in less accessible chromatin may be a key feature of the evolutionary mechanism of segment diversification.

Overall, our studies indicate the role played by chromatin accessibility in Hox target selection and the observation that Hox binding is much more closely correlated with chromatin accessibility than with binding affinity has implications for other systems in understanding the relationship between genome sequence and transcription factor binding. It fits with studies on transcription factor binding in the *Drosophila* blastoderm [4,42] and echoes the recent observation that the interpretation of the gradient of the homeodomain protein Bicoid, in establishing the anterior-posterior axis in the *Drosophila* embryo, is more dependent on chromatin accessibility than on the binding affinity of target sites [43].

## Materials and Methods

### Cell culture

Kc167 cells (obtained from the Drosophila Genomics Resource Center) were cultured in Schneider’s medium supplemented with 5% fetal calf serum and antibiotics at 25°C.

### Expression plasmid cloning

Coding sequences (CDSs) for the eGFP-tagged Hox proteins Ubx, Abd-A and Abd-B, and for the Hth cofactor derived from Hox-vectors produced by Beh et al. [18]. CDSs for the remaining Hox proteins Lab, Pb, Dfd, Scr, Antp and for Exd and Gcm were amplified from a cDNA preparation (QIAGEN, 205310) of 0-12 hour old embryos via nested-PCR, starting with primers specific to flanking UTRs of each target CDS. All DNA amplifications were done using a Phusion High-Fidelity DNA Polymerase (NEB, M0530).

For transient transfection, sequences encoding eGFP-tagged transcription factors were cloned into the pMT expression vector (Invitrogen V4120-20), which employs the inducible *Drosophila* metallothionein promoter to drive transgene expression using a suitable CuSO_4_ concentration in the growing medium.

To generate Kc167 cell lines stably carrying inducible eGFP-tagged factors (stable lines), CDSs were cloned into the pMT-puro expression vector (Addgene, #17923), which uses a puromycin selection system. We produced stable lines by selecting cells in medium with 5 µg/ml of puromycin after transfection with pMT-puro-Hox constructs (see below).

We produced vectors expressing either single eGFP-tagged Hox proteins, Hth and Gcm (monocistronic vectors), or eGFP-tagged Hox factors in association with Hth, eGFP-Exd in association with Hth, and specific Hox factors in association with Gcm (bicistronic vectors). We employed the T2A peptide self-cleavage system for multicistronic constructs. All constructs were sequence-verified.

### Transfection

Transient transfection was performed according to Beh et al. [18]. Briefly, Kc167 cells harvested in log phase were used to seed 10 cm dishes (Corning Inc. 353003) at a density of 2.5 × 107 cells per dish and transfection was performed using FuGENE 6 Transfection Reagent (Promega E2691) according to the manufacturer’s instructions. Dishes were then incubated at 25°C for approximately 14 hours. For stable transfection, 2 × 106 cells were re-suspended in 10 ml Schneider’s medium containing the transfection solution (70µl OPTIMEM, 3µl Fugene and 2µg plasmid DNA) and seeded into 10 cm dishes. After incubation at 25°C for approximately 18 hours, the medium was replaced with standard Schneider’s and cells were cultured for approximately 24 hours before starting puromycin selection.

### Induction of gene expression, fixation and FACS

For transiently transfected cells, medium was replaced by 10 ml of Schneider’s medium/1 mM CuSO_4_ and dishes were incubated at 25°C for 4 hours to induce GFP-Hox expression. In the case of stable lines, CuSO_4_ concentrations and induction times varied between lines and were adjusted to provide optimal expression levels prior to FACS sorting. Cell fixation and FACS sorting methods were as in Beh et al. [18]. Cells destined for ATAC experiments were not fixed and were FACS sorted into PBS, 0.1% BSA instead of PBS, 0.01% Triton X-100. An equal number of cells (106) was sorted for all samples.

### ChIP and ChIP-seq library preparation

ChIP was performed as in Beh et al. [18] except the anti-GFP antibody used in this study was from Sigma (G1544; 2µl per ChIP). ChIP and input DNA were re-suspended in 20µl of TE buffer. 10 µl of ChIP DNAs and 400pg of input DNA in 10 µl TE buffer were used to produce sequencing libraries using the SMARTer ThruPLEX DNA-seq Kit (Takara Bio Inc.) in accordance with the sample preparation guide. Fourteen cycles of amplification were used for all libraries.

ATAC-seq

ATAC-seq libraries were prepared according to Buenrostro et al. [44]. Final libraries were size selected to contain molecules of 150-700 bp using AMPure XP beads (Beckman Coulter).

### Sequencing and data processing

Libraries were either sequenced on the Illumina HiSeq 2000 or HiSeq 4000 platforms at the CRUK Cambridge Institute Genomics Core. ChIP-seq and ATAC-seq reads were aligned to the Drosophila melanogaster genome BDGP release 6 (dm6) excluding scaffolds using bowtie (v 1.2.2) with the -m1 option. Reads were then converted to bam files with Samtools (v 1.3.1). ChIP-seq peak detection for each biological replicate using the input as background was performed with MACS2 (v 2.1.1.20160309) using --keep-dup 1, --call-summits and -q 1e-2 and -q 1e-10 options. Binding regions overlapping exon regions contained in the plasmid were then removed. Bound regions were defined as the union of overlapping regions detected by MACS2 across both replicates at a given stringency. Unless stated otherwise we use q-value 1e-2 in the figure plots.

ATAC-seq reads aligning to the + strand were offset by +4 bp, and reads aligning to the - strand were offset −5 bp to represent the centre of the transposase binding, then the reads were extended by 5bp on either side. Open regions for the basal Kc-cells were then called using MACS2 with options --shift −45 --extsize 100 and -q 1e-2 for each of the 3 replicates. Basal ATAC core open regions were defined as the union of open regions present in at least 2 of the replicates. See Tables S1-3 for ChIP-Seq read overview, ChIP-Seq binding region numbers and ATAC-seq read overview. The ChIP-Seq and ATAC-Seq data are available from GEO under accession number GSE122575.

### Hox group peak regions

To define bound regions between all Hox and Hox+Cofactor(s) ChIP samples the sub-peak summit positions at MACS q-value 1e-10 were grouped using GenomicRanges R package [45]. Starting with the sample with the largest number of sub-peak summits these summit positions were extended +/-100 bp and then overlapped with the extended summits of the next sample. A new centre position was then calculated using the mean position between all sub-peak summits belonging in this grouped region. All non-overlapping summit positions were taken to the next round. Finally, group regions containing less than 2 members were removed. This resulted in 200 bp peak regions. For the transient transfected data which includes all 8 Hox and Hox+Hth samples this resulted in 15,945 regions, called Hox group peaks. For the stable cell line data we used the 3 Hox samples (Dfd, Ubx and Abd-B) and Hox+cofactors (Hth and Gcm) samples which resulted in 21,002 regions now called Hox group peaks - stable. Each Hox group peak region was then flagged as bound by a specific Hox (or Hox+cofactor) if peak regions of both replicates at the selected stringency overlapped (we used min overlap 1 bp throughout). Additionally the regions were flagged as open if they overlapped with the Kc-cell basal core open regions. The Hox group peak regions are detailed in Table S4.

### Cofactor-enhanced binding analysis

Reads overlapping the Hox group peaks were counted using the union method of the summarizeOverlaps function in the GenomicAlignments R package by extending the reads by their fragment size (as determined by MACS2). The count table was then processed with edgeR R package [46] as follows: reads were normalised using the loess method (as per csaw R package; [47]) to remove trended bias, then the dispersions were calculated and the glmQLFit function used to fit a quasi-likelihood negative binomial generalized log-linear model to count data. Differential binding (DB) analysis was performed per pair-wise comparison between two samples using a threshold of fdr <= 0.01 and logFC >= 1 (in this case log difference of binding signal), additionally both replicates of the DB sample were required to be bound at macs q-value 1e-2 (Table S5).

### ChIP and ATAC scores

The ChIP-seq reads of both replicates were extended to match the mean fragment size. ATAC-seq reads of both replicates were extended by 100 bp centered on the Tn5 cut-site. Bedgraph files were then created using MACS2 pileup and scaled to reads-per-million, counting reads overlapping the Hox group peaks for each experiment. The profiles were then binned at 20 bp resolution using the mean score. ChIP or ATAC scores of selected regions were then calculated as the mean profile score of overlapping bins.

### Venn Diagrams

The highest binding score position in regions bound by both replicates at the selected stringency was extended by +/- 200 bp. To deal with the problem of one region overlapping with two (or more) regions in the other sample we created the union of these regions across the three Hox samples under investigation, thus creating a unique region set. For each individual Hox the overlap with the union region was quantified and plotted as a proportion sized Venn diagram using the eulerr R package [48].

### Motif analysis

Motif enrichment analysis was performed using PWMEnrich R package [49]. Motif enrichment scores [log_10_(1/p-value)] were grouped by transcription factor and individual motifs plotted as dot plots with the median as coloured bar, or grouped into HoxA* (Lab, Pb, Dfd, Scr, Antp, Ubx, Abd-A) and Abd-B and plotted as boxplots using R.

Jaspar Hox PWMs were truncated to 7-mers for Total Binding Affinity (TBA; [20]), Hox site counting and maxScore analysis. We then combined the PWMs of Lab, Pb, Dfd, Scr, Antp, Ubx, Abd-A to a new PWM HoxA and renamed Abd-B to HoxB.

TBA was calculated across the Hox group peaks (Fig 3D, Fig 4D,E, Fig S1C,D) or binding summit regions extended by +/- 100bp (Fig 3B) using MatrixRider R package [50]. Sequences were searched using the truncated Hox 7-mer PWMs with Biostrings R package [51], matchPWM function with min.score=80% on both strands and all possible sites (allowing overlaps) counted. Additionally the highest score within each sequence for each PWM was extracted (using min.score>=50%).

De-novo motif discovery was performed using HOMER [52] on the cofactor-enhanced binding regions (Hox+Hth) (Table S5, Fig S4). All sequence logos were plotted using seqLogo R package [53].

Consensus matrixes for Fig 7A were created with Biostrings R package finding all matches to TGATTDAT (where D=A or G or T), based on the in vitro SELEX Exd-Hox sites [16] and our HOMER de-novo motifs, allowing 1 mismatch in cofactor-enhanced binding regions. The binding regions used were: Class 1 unique Exd/Hth enhanced Lab bound; Class 2 unique Exd/Hth enhanced Pb or Dfd or Scr bound and Class 3 unique Exd/Hth enhanced Antp or Ubx or Abd-A or Abd-B bound.

The top 15 prevalent 8-mer sequence patterns in Exd/Hth enhanced binding regions were determined using Biostrings, masking identified kmers after each round.

SELEX raw data was downloaded from GSE65073 [54] and reprocessed using the SELEX R package [55] with optimal length = 12 and markov order = 5, to obtain complete affinity tables for each Exd-Hox experiment. The affinities for the three Exd-Hox class patterns in Fig 7A were then looked up locating any 12-mer containing these patterns (or the reverse complement) and plotted as a stripchart plot using R.

### Correlation Heatmap

The union of all regions bound by Hox or Hox+Hth at MACS q-value 1e-2 was tiled into 20 bp windows and reads overlapping each window were counted using csaw R package [47]. Reads were then normalised by library size and transformed to counts per million. The correlation between the samples was then plotted using heatmap.2 from the gplots R package.

### Chromatin accessibility analysis

The 10bp adjusted ATAC-seq reads overlapping the 21,002 Hox group peaks - stable regions were counted as above. These counts were then processed as for the cofactor-enhanced binding analysis. We defined significantly increased chromatin accessibility regions as edgeR fdr <= 0.01 and logFC >= 1.5 comparing induced versus non-induced samples (Table S6).

### Occupancy heatmaps in open regions

Open chromatin regions in basal Kc-cells (16,118 regions ranging in size between 100 bp – 2413 bp) were tiled into 200 bp bins as follows: smaller regions were resized to 200 bp fixed on the center of each region and larger regions were split into 200 bp tiles. The tiles were then classified as bound or not bound if they overlapped a Hox bound region. Mean ATAC scores of basal Kc-cells and TBA for HoxA or HoxB PWMs were calculated per tile. The log of these scores was then linearly binned into 40 bins and a heatmap plotted. For the “All regions” plot the heatmap colours shows the location of highest density of these tiles. The colours in the other plots represent the proportion of Hox bound within each bin. We then assessed the correlation R2 of occupancy (proportion bound per bin) with chromatin accessibility (ATAC scores) or binding affinity (TBA), shown as scatterplots.

## Data Availability

All ChIP-Seq and ATAC-Seq data files are available from the GEO database (accession number GSE122575).

Supplementary Tables are available at https://doi.org/10.5281/zenodo.1491604

## Author Contributions and Notes

DP and BF performed the research and analysed the data, RW and SR conceived the project and analysed the data all authors wrote the paper. The authors declare no conflict of interest. This work is supported by a grant to RW and SR from the Biotechnology and Biological Sciences Research Council (BBSRC; www.bbsrc.ac.uk); Grant No. BB/M007081/1. The funders had no role in study design, data collection and analysis, decision to publish, or preparation of the manuscript.

## Acknowledgments

We thank Hanno Fischer for the fly image in Figure 1.

## Supplementary Figures

**S1 Figure.**
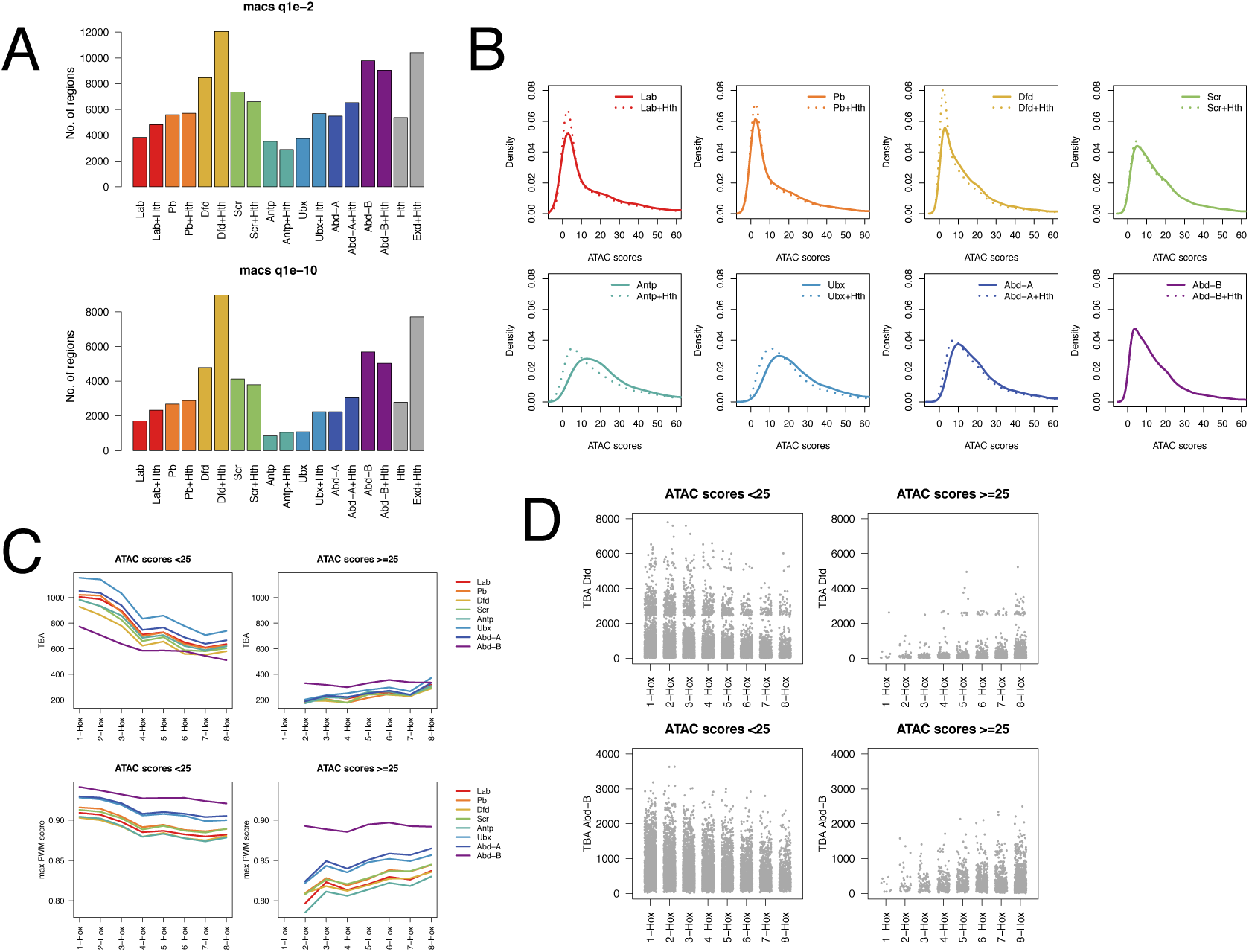
Binding region numbers, chromatin accessibility profiles and the association between Hox specificity and binding site affinity. A: Numbers of regions bound by Hox and Hox+Hth for both replicates at q-value 1e-2 (upper) and higher stringency q-value 1e-10 (lower). B: Density plots of mean ATAC-seq scores for regions bound by Hox proteins with and without Exd/Hth showing the effect of the cofactors on the chromatin accessibility profile. Solid lines: Hox alone, dotted lines: Hox in presence of Exd/Hth. C: Plots showing relationship between Hox selectivity and either (top row) TBA or (bottom row) highest PWM score for binding regions as in Figure 4A, but separated according to chromatin accessibility with more closed on left (ATAC scores < 25) and more open on right (ATAC scores >= 25). TBA is shown for each jaspar 7-mer Hox PWM. The maxPWM score is the mean of the highest PWM score per region. Whilst the more closed regions show a positive association between Hox selectivity and both TBA and maxPWM score, the trend is reversed for the more open regions. Number of regions with ATAC scores < 25 is 13507 and number of regions with ATAC scores >= 25 is 2448. Data for 1-Hox is omitted due to low number of regions in this bin. D: TBA data for Dfd and Abd-B PWMs as in (C) plotted as strip plots showing clear reversal of association between TBA and Hox selectivity in more open versus less open chromatin.

**S2 Figure.**
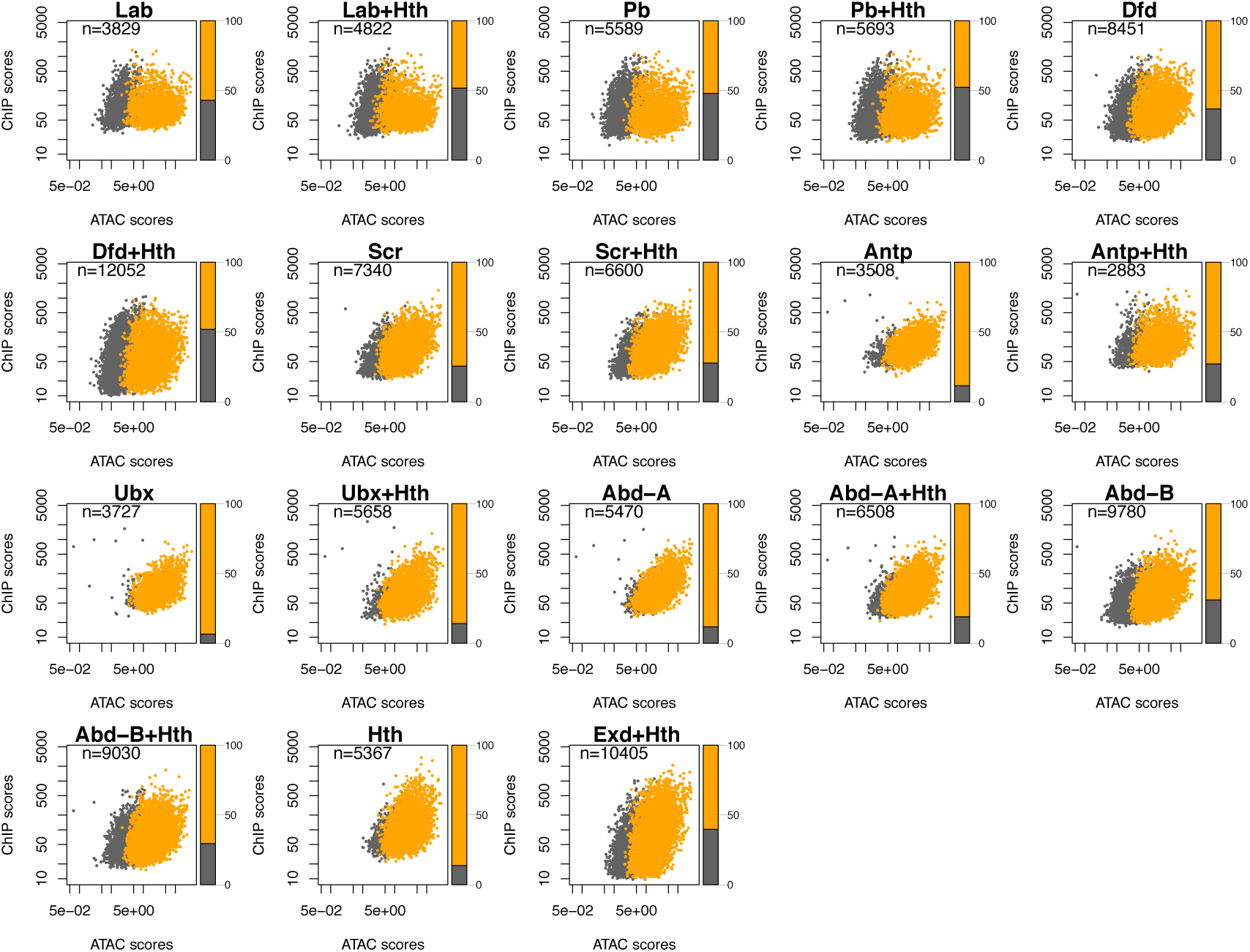
Hox, Hox+Hth, Hth and Exd binding in open and closed chromatin. Scatter plots of ChIP score versus chromatin accessibility for each Hox and Hox+Hth ChIP and for Hth ChIP and ChIP of Exd in the presence of Hth. The binding summits positions bound at q-value 1e-10 were extended +/- 100bp and the mean of the ChIP scores and the mean of the basal Kc-cell ATAC scores plotted. Regions which overlap with open chromatin regions as determined using the same ATAC-seq data using macs2 at q1e-2 are shown in orange, grey regions indicate closed chromatin, regions do not overlap chromatin called as open. The barplot at the side indicate the proportion of closed and open regions. The plots show the different degrees of binding to closed chromatin for different Hox proteins and the general increase in binding to closed chromatin in the presence of Exd/Hth. Hth shows little binding to closed chromatin. Exd apparently shows more binding in closed chromatin, but note that much of this effect is due to the inclusion of lower ChIP scoring regions in the Exd+Hth plot. This is a result of lower scoring peaks reaching significance due to the low background in this ChIP.

**S3 Figure.**
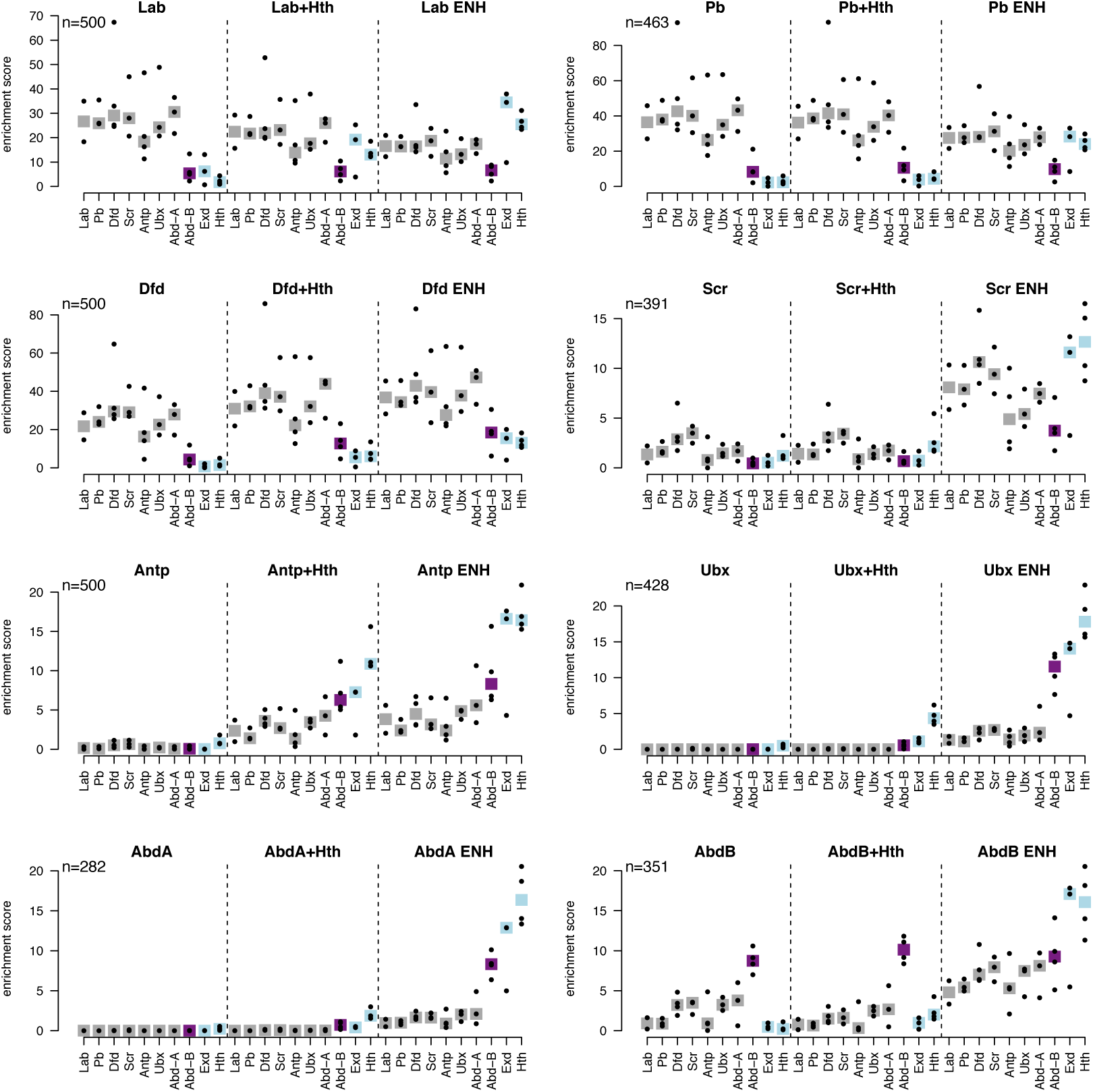
Motif analysis showing individual motifs. Motif analysis comparing motif enrichment for Hox group regions for Hox alone (Hox), Hox in the presence of Exd/Hth (Hox+Hth) and Exd/Hth cofactor enhanced binding regions (Hox ENH) using the top (highest ChIP score) 500 regions from each class (or matched numbers to the ENH set where less than 500). Plot titles indicate binding region set used and motifs are indicated on the x-axis. Enrichment analysis was performed using PWMEnrich for the Hox motifs in the MotifDb database. Enrichment scores [log_10_(1/p-value)] for individual motifs are indicated (dots) together with the median for each motif set (bar). Lab, Pb, Dfd, Scr, Antp, Ubx and Abd-A (grey bar), Abd-B (purple bar), and the Exd and Hth motifs (light blue). Note the differences in Y-axis scale. Generally the provision of Exd/Hth has little effect on the Hox motif enrichments although the Exd and Hth motifs show increased enrichment. For Antp, Ubx and Abd-A the provision of Exd/Hth increases the relative enrichment of the Abd-B motif relative to the other Hox motifs, which may reflect the in vitro defined phenomenon of latent specificity [16].

**S4 Figure.**
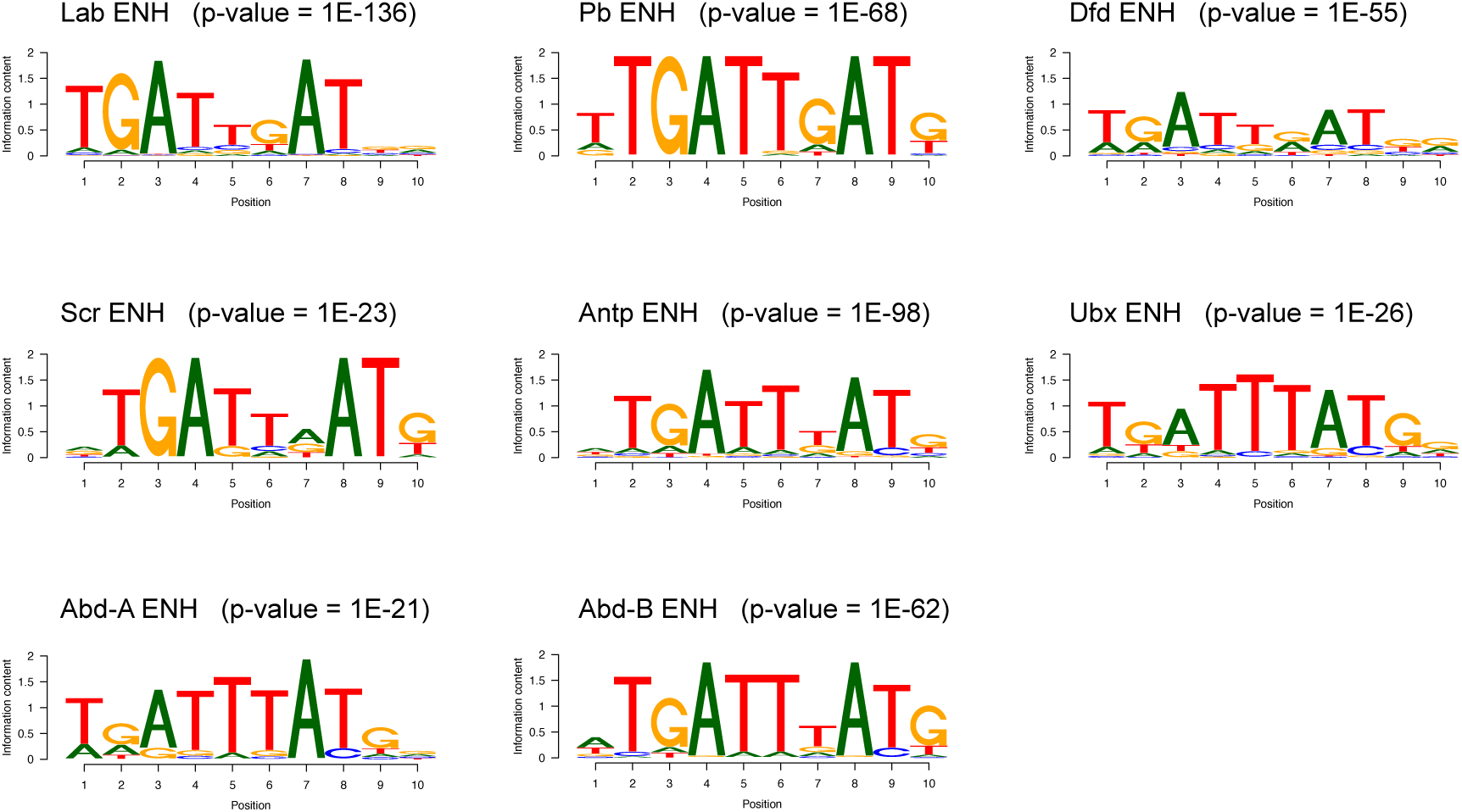
De-novo motif analysis of Exd/Hth cofactor enhanced binding regions. PWMs for enriched motifs in Exd/Hth cofactor enhanced binding regions (Hox ENH) using HOMER. Enrichment p-values are shown.

**S5 Figure.**
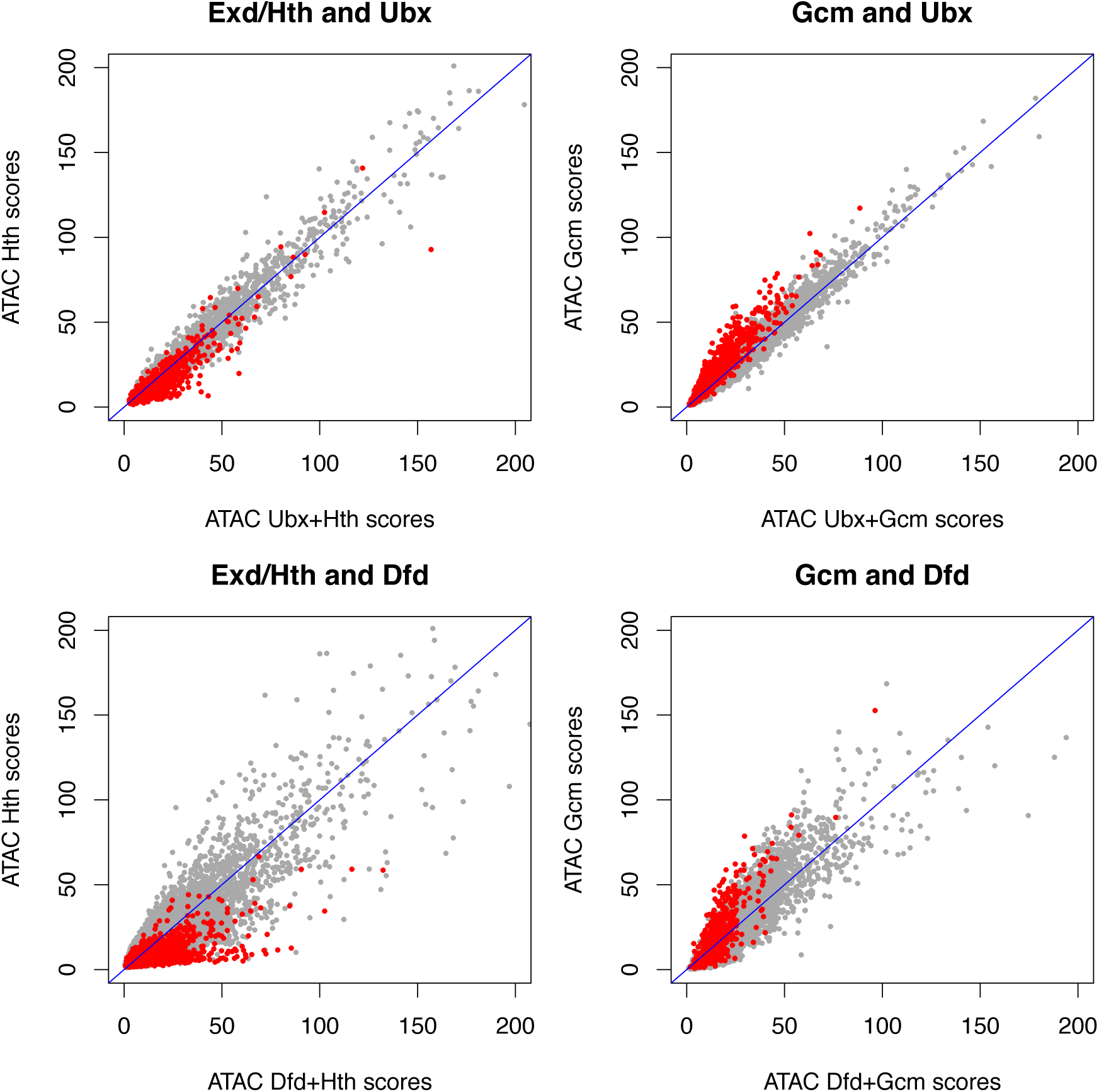
Comparing the effects of Exd/Hth and Gcm: Chromatin accessibility in Hox + Exd/Hth compared to Hox + Gcm. Scatterplots comparing the effect on chromatin accessibility of providing Exd/Hth versus Gcm in addition to Hox proteins (Ubx or Dfd). Mean ATAC score per binding region are plotted for all the bound regions in Hox+Hth or Hox+Gcm respectively. On the Y-axis the ATAC scores are plotted for these regions in Hth or Gcm alone and on the X-axis the ATAC scores are plotted with the addition of Hox proteins (Ubx or Dfd). The Exd/Hth-enhanced or Gcm-enhanced regions respectively are shown in red. In the case of Exd/Hth, the Exd/Hth-enhanced Hox binding regions predominantly lie below the diagonal; i.e. they have increased chromatin accessibility in Hox+Hth than in Hth alone. This is not the case for Gcm. This illustrates the difference in the relationship between Hox proteins and Exd/Hth compared to that between Hox proteins and Gcm. In the case of Exd/Hth the Hox proteins collaborate in promoting chromatin accessibility, whereas in the case of Gcm, the chromatin accessibility state is predominantly controlled by Gcm.

**S6 Figure.**
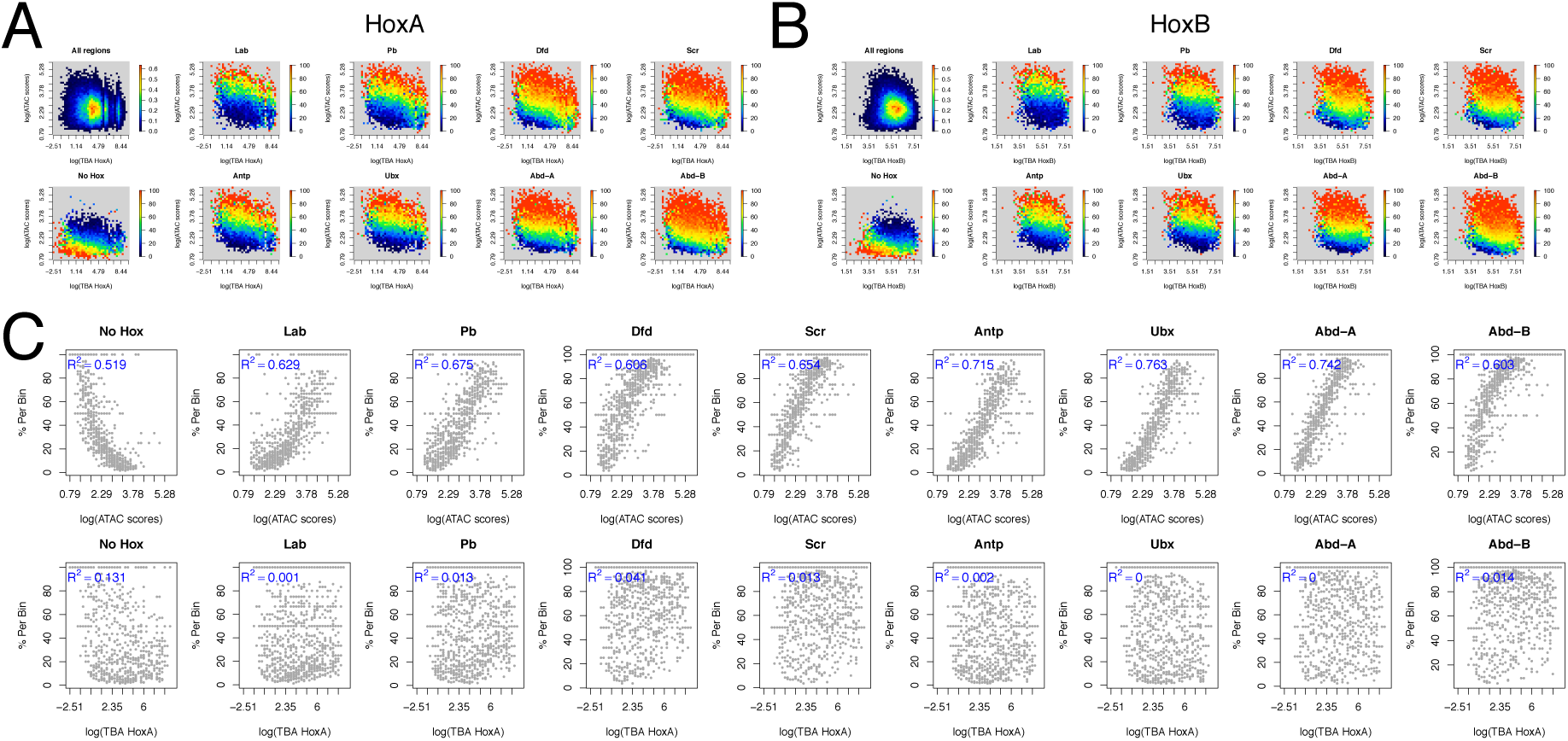
Hox occupancy is more strongly associated with binding region chromatin accessibility than with binding affinity. Scatter plots of chromatin accessibility (log[ATAC scores]) versus binding affinity (log[TBA HoxA]) for chromatin regions classified as “open”. Open chromatin regions were divided into 200bp tiles and the mean ATAC score and TBA for HoxA PWM (A) or HoxB PWM (B) calculated per tile. The log of these scores was then linearly binned into 40 bins on each axis. For the “All regions” plot the heatmap shows the density distribution. For the other plots, the heatmap shows the percentage of tiles bound by the specified Hox protein per bin or for “No Hox” the percentage of tiles not bound by any Hox protein. (C) Scatter plots show the strong correlation of occupancy (% per bin) with chromatin accessibility (log[ATAC scores]; upper row) and the poor correlation with binding affinity ((log[TBA HoxA]; lower row). Data as in (A).

## Supplementary tables available at https://doi.org/10.5281/zenodo.1491604

**Table S1.** ChIP-seq read overview.

| Transient Transfections |  |  |  |  | Stable Lines |  |  |  |
| --- | --- | --- | --- | --- | --- | --- | --- | --- |
| Sample | Total_reads | Mapped_reads | Platform |  | Sample | Total_reads | Mapped_reads | Platform |
| Lab_225 | 35,652,497 | 19,776,331 | HiSeq 4000 |  | Dfd_247 | 32,843,529 | 18,779,761 | HiSeq 4000 |
| Lab_226 | 31,656,390 | 17,331,670 | HiSeq 4000 |  | Dfd_248 | 17,095,829 | 10,292,158 | HiSeq 4000 |
| Lab_input | 21,993,989 | 12,275,906 | HiSeq 4000 |  | Dfd_input | 27,181,802 | 16,427,314 | HiSeq 4000 |
| Lab+Hth_228 | 19,535,480 | 10,340,548 | HiSeq 4000 |  | Dfd+Hth_243 | 21,869,632 | 14,114,182 | HiSeq 4000 |
| Lab+Hth_229 | 32,379,953 | 18,106,788 | HiSeq 4000 |  | Dfd+Hth_244 | 25,725,965 | 16,369,234 | HiSeq 4000 |
| Lab+Hth_input | 19,029,087 | 10,947,693 | HiSeq 4000 |  | Dfd+Hth_input | 23,240,584 | 14,492,712 | HiSeq 4000 |
| Pb_201 | 24,903,026 | 12,045,654 | HiSeq 4000 |  | Dfd+Gcm_249 | 36,208,630 | 21,210,268 | HiSeq 4000 |
| Pb_202 | 24,425,098 | 11,872,562 | HiSeq 4000 |  | Dfd+Gcm_250 | 20,584,442 | 12,196,295 | HiSeq 4000 |
| Pb_input | 25,449,790 | 14,884,704 | HiSeq 4000 |  | Dfd+Gcm_input | 41,719,566 | 25,980,366 | HiSeq 4000 |
| Pb+Hth_203 | 32,955,638 | 13,326,705 | HiSeq 4000 |  | Ubx_217 | 40,574,216 | 22,651,116 | HiSeq 4000 |
| Pb+Hth_204 | 16,736,809 | 6,084,266 | HiSeq 4000 |  | Ubx_218 | 33,836,429 | 19,965,122 | HiSeq 4000 |
| Pb+Hth_input | 32,830,552 | 19,085,144 | HiSeq 4000 |  | Ubx_input | 23,514,868 | 14,304,826 | HiSeq 4000 |
| Dfd_113 | 20,422,023 | 11,993,655 | HiSeq 4000 |  | Ubx+Hth_213 | 37,082,776 | 21,381,969 | HiSeq 4000 |
| Dfd_114 | 18,352,596 | 11,186,142 | HiSeq 4000 |  | Ubx+Hth_214 | 32,117,571 | 18,761,644 | HiSeq 4000 |
| Dfd_input | 43,905,501 | 27,275,300 | HiSeq 4000 |  | Ubx+Hth_input | 26,843,880 | 16,594,979 | HiSeq 4000 |
| Dfd+Hth_117 | 35,673,391 | 22,902,065 | HiSeq 4000 |  | Ubx+Gcm_235 | 45,711,964 | 26,450,967 | HiSeq 4000 |
| Dfd+Hth_118 | 31,982,490 | 20,279,888 | HiSeq 4000 |  | Ubx+Gcm_236 | 44,458,658 | 26,238,573 | HiSeq 4000 |
| Dfd+Hth_input | 54,438,852 | 33,037,109 | HiSeq 4000 |  | Ubx+Gcm_input | 41,935,371 | 24,920,987 | HiSeq 4000 |
| Scr_115 | 26,549,969 | 15,425,586 | HiSeq 4000 |  | Abd-B_239 | 39,832,402 | 26,838,262 | HiSeq 4000 |
| Scr_116 | 25,966,588 | 14,805,387 | HiSeq 4000 |  | Abd-B_240 | 72,131,533 | 45,950,537 | HiSeq 4000 |
| Scr_input | 45,557,965 | 26,756,096 | HiSeq 4000 |  | Abd-B_input | 37,184,024 | 23,188,692 | HiSeq 4000 |
| Scr+Hth_119 | 20,753,220 | 12,635,134 | HiSeq 4000 |  | Abd-B+Hth_241 | 33,136,444 | 22,093,185 | HiSeq 4000 |
| Scr+Hth_120 | 23,495,495 | 14,361,088 | HiSeq 4000 |  | Abd-B+Hth_242 | 22,183,688 | 14,712,517 | HiSeq 4000 |
| Scr+Hth_input | 31,207,186 | 18,740,677 | HiSeq 4000 |  | Abd-B+Hth_input | 34,244,185 | 21,439,677 | HiSeq 4000 |
| Antp_67 | 33,839,220 | 15,204,772 | HiSeq 4000 |  | Gcm_219 | 38,041,560 | 23,403,544 | HiSeq 4000 |
| Antp_68 | 42,079,289 | 18,791,811 | HiSeq 4000 |  | Gcm_220 | 38,754,938 | 23,227,820 | HiSeq 4000 |
| Antp_input | 39,031,841 | 22,009,066 | HiSeq 4000 |  | Gcm_input | 39,285,389 | 23,608,832 | HiSeq 4000 |
| Antp+Hth_75 | 28,360,324 | 12,993,786 | HiSeq 4000 |  | Hth_215 | 42,237,064 | 26,788,395 | HiSeq 4000 |
| Antp+Hth_76 | 29,674,613 | 13,895,182 | HiSeq 4000 |  | Hth_216 | 36,857,352 | 23,428,980 | HiSeq 4000 |
| Antp+Hth_input | 21,860,819 | 13,107,646 | HiSeq 4000 |  | Hth_input | 24,560,848 | 15,060,156 | HiSeq 4000 |
| Ubx_1 | 26,255,388 | 12,094,751 | HiSeq 2500 |  |  |  |  |  |
| Ubx_2 | 27,539,748 | 13,518,594 | HiSeq 2500 |  |  |  |  |  |
| Ubx_input | 48,617,397 | 27,131,432 | HiSeq 2500 |  |  |  |  |  |
| Ubx+Hth_1 | 25,473,053 | 11,488,879 | HiSeq 2500 |  |  |  |  |  |
| Ubx+Hth_2 | 36,697,607 | 16,089,206 | HiSeq 2500 |  |  |  |  |  |
| Ubx+Hth_input | 25,155,474 | 14,512,496 | HiSeq 2500 |  |  |  |  |  |
| Abd-A_49 | 18,734,401 | 10,471,829 | HiSeq 2500 |  |  |  |  |  |
| Abd-A_50 | 23,927,894 | 12,894,433 | HiSeq 2500 |  |  |  |  |  |
| Abd-A_input | 21,996,208 | 12,532,916 | HiSeq 2500 |  |  |  |  |  |
| Abd-A+Hth_53 | 23,305,535 | 12,221,375 | HiSeq 2500 |  |  |  |  |  |
| Abd-A+Hth_54 | 16,228,122 | 8,881,775 | HiSeq 2500 |  |  |  |  |  |
| Abd-A_input | 34,576,407 | 20,530,123 | HiSeq 2500 |  |  |  |  |  |
| Abd-B_51 | 22,788,335 | 12,470,631 | HiSeq 2500 |  |  |  |  |  |
| Abd-B_52 | 23,583,829 | 13,700,974 | HiSeq 2500 |  |  |  |  |  |
| Abd-B_input | 21,996,208 | 12,532,916 | HiSeq 2500 |  |  |  |  |  |
| Abd-B+Hth_55 | 22,523,431 | 13,915,019 | HiSeq 2500 |  |  |  |  |  |
| Abd-B+Hth_56 | 22,403,332 | 7,508,223 | HiSeq 2500 |  |  |  |  |  |
| Abd-B_input | 34,576,407 | 20,530,123 | HiSeq 2500 |  |  |  |  |  |
| Hth_209 | 38,180,841 | 21,305,298 | HiSeq 4000 |  |  |  |  |  |
| Hth_210 | 37,150,247 | 20,663,723 | HiSeq 4000 |  |  |  |  |  |
| Hth_input | 29,208,741 | 16,401,950 | HiSeq 4000 |  |  |  |  |  |
| Exd+Hth_232 | 28,385,162 | 17,462,671 | HiSeq 4000 |  |  |  |  |  |
| Exd+Hth_233 | 23,369,536 | 14,523,255 | HiSeq 4000 |  |  |  |  |  |
| Exd+Hth_input | 32,952,245 | 18,571,046 | HiSeq 4000 |  |  |  |  |  |

**Table S2.** ChIP-Seq binding region numbers. Union gives the number of binding regions present in both replicates

**Transient transfections**
| **Sample** | **q1e-2** |  |  | **q1e-10** |  |  |
| --- | --- | --- | --- | --- | --- | --- |
|  | **rep1** | **rep2** | **union** | **rep1** | **rep2** | **union** |
| **Lab** | 4206 | 4742 | **3835** | 1790 | 2089 | **1704** |
| **Lab+Hth** | 5406 | 5922 | **4827** | 2493 | 2741 | **2325** |
| **Pb** | 7204 | 6519 | **5598** | 3301 | 2903 | **2682** |
| **Pb+Hth** | 7605 | 6204 | **5708** | 3887 | 2990 | **2883** |
| **Dfd** | 9773 | 9365 | **8467** | 5703 | 5102 | **4782** |
| **Dfd+Hth** | 13195 | 12952 | **12054** | 9731 | 9510 | **8958** |
| **Scr** | 8488 | 8576 | **7355** | 4700 | 4793 | **4127** |
| **Scr+Hth** | 7616 | 8581 | **6603** | 4257 | 5285 | **3797** |
| **Antp** | 3970 | 5468 | **3523** | 921 | 1451 | **846** |
| **Antp+Hth** | 3289 | 4728 | **2887** | 1154 | 1704 | **1043** |
| **Ubx** | 5271 | 4372 | **3752** | 1581 | 1209 | **1083** |
| **Ubx+Hth** | 6473 | 7619 | **5684** | 2428 | 3112 | **2242** |
| **Abd-A** | 6251 | 7242 | **5487** | 2412 | 3203 | **2234** |
| **Abd-A+Hth** | 7525 | 7867 | **6528** | 3416 | 3617 | **3040** |
| **Abd-B** | 11414 | 10906 | **9786** | 6915 | 6133 | **5685** |
| **Abd-B+Hth** | 10976 | 10144 | **9044** | 6444 | 5476 | **5027** |
| **Hth** | 6572 | 5872 | **5373** | 3551 | 2921 | **2781** |
| **Exd+Hth** | 11451 | 11590 | **10409** | 8623 | 8383 | **7695** |

**Stable cell lines**
| **Sample** | **q1e-2** |  |  | **q1e-10** |  |  |
| --- | --- | --- | --- | --- | --- | --- |
|  | **rep1** | **rep2** | **union** | **rep1** | **rep2** | **union** |
| **Exd+Hth_stable** | 11451 | 11590 | **10409** | 8623 | 8383 | **7695** |
| **Gcm_stable** | 12301 | 11863 | **10765** | 6683 | 6125 | **5831** |
| **Hth_stable** | 9891 | 10045 | **8999** | 6713 | 7238 | **6225** |
| **Dfd_stable** | 12332 | 11500 | **10380** | 6261 | 5566 | **5145** |
| **Dfd+Hth_stable** | 11156 | 11699 | **9867** | 6464 | 7260 | **5981** |
| **Dfd+Gcm_stable** | 10261 | 10370 | **8893** | 4533 | 4506 | **3924** |
| **Ubx_stable** | 6251 | 6019 | **5281** | 2206 | 2066 | **1833** |
| **Ubx+Hth_stable** | 5406 | 6852 | **4885** | 2415 | 3422 | **2269** |
| **Ubx+Gcm_stable** | 10035 | 11039 | **8846** | 4612 | 6140 | **4275** |
| **Abd-B_stable** | 14696 | 14839 | **13511** | 11371 | 11222 | **10375** |
| **Abd-B+Hth_stable** | 15441 | 14902 | **13741** | 11152 | 10050 | **9418** |

**Table S3.** Stable cell lines ATAC-seq read overview. # ATAC_Kc represent standard Kc167 cells, all other samples are stable Kc-cell lines containing pMT-puro-Hox plasmids # i = induced (CuSO4), n = non-induced

| Sample | Total_reads | Mapped_reads | Platform |
| --- | --- | --- | --- |
| ATAC_Kc_7 | 112,702,038 | 90,189,400 | HiSeq 4000 |
| ATAC_Kc_10 | 113,325,609 | 87,818,612 | HiSeq 4000 |
| ATAC_Kc_11 | 131,356,618 | 102,734,912 | HiSeq 4000 |
| ATAC_Dfd_i_114 | 33,979,005 | 26,174,422 | HiSeq 4000 |
| ATAC_Dfd_i_115 | 47,498,366 | 35,298,598 | HiSeq 4000 |
| ATAC_Dfd_n_116 | 8,106,401 | 5,926,680 | HiSeq 4000 |
| ATAC_Dfd_n_117 | 16,827,993 | 12,395,692 | HiSeq 4000 |
| ATAC_Dfd+Hth_i_106 | 20,645,066 | 15,412,147 | HiSeq 4000 |
| ATAC_Dfd+Hth_i_107 | 33,393,909 | 25,018,382 | HiSeq 4000 |
| ATAC_Dfd+Hth_n_108 | 45,517,692 | 33,527,834 | HiSeq 4000 |
| ATAC_Dfd+Hth_n_109 | 36,571,843 | 27,512,683 | HiSeq 4000 |
| ATAC_Dfd+Gcm_i_118 | 19,123,450 | 14,542,498 | HiSeq 4000 |
| ATAC_Dfd+Gcm_i_119 | 31,521,343 | 23,755,454 | HiSeq 4000 |
| ATAC_Dfd+Gcm_n_120 | 32,482,773 | 23,479,769 | HiSeq 4000 |
| ATAC_Dfd+Gcm_n_121 | 22,607,002 | 16,957,603 | HiSeq 4000 |
| ATAC_Ubx_i_62 | 33,125,682 | 23,727,431 | HiSeq 4000 |
| ATAC_Ubx_i_63 | 47,570,852 | 34,116,016 | HiSeq 4000 |
| ATAC_Ubx_n_64 | 13,589,247 | 10,291,131 | HiSeq 4000 |
| ATAC_Ubx_n_65 | 24,444,782 | 18,040,490 | HiSeq 4000 |
| ATAC_Ubx+Hth_i_54 | 34,488,251 | 24,207,832 | HiSeq 4000 |
| ATAC_Ubx+Hth_i_55 | 30,566,866 | 23,181,127 | HiSeq 4000 |
| ATAC_Ubx+Hth_n_56 | 50,532,615 | 38,846,046 | HiSeq 4000 |
| ATAC_Ubx+Hth_n_57 | 33,874,296 | 25,808,470 | HiSeq 4000 |
| ATAC_Ubx+Gcm_i_90 | 47,304,001 | 37,298,997 | HiSeq 4000 |
| ATAC_Ubx+Gcm_i_91 | 52,695,324 | 43,620,118 | HiSeq 4000 |
| ATAC_Ubx+Gcm_n_92 | 30,341,723 | 22,678,298 | HiSeq 4000 |
| ATAC_Ubx+Gcm_n_93 | 13,620,559 | 10,014,547 | HiSeq 4000 |
| ATAC_Abd-B_i_98 | 22,418,822 | 17,167,407 | HiSeq 4000 |
| ATAC_Abd-B_i_99 | 24,413,883 | 19,340,938 | HiSeq 4000 |
| ATAC_Abd-B_n_100 | 20,580,779 | 15,982,847 | HiSeq 4000 |
| ATAC_Abd-B_n_101 | 27,866,139 | 21,673,485 | HiSeq 4000 |
| ATAC_Abd-B+Hth_i_102 | 41,645,070 | 34,657,116 | HiSeq 4000 |
| ATAC_Abd-B+Hth_i_103 | 23,886,047 | 18,624,131 | HiSeq 4000 |
| ATAC_Abd-B+Hth_n_104 | 19,484,964 | 14,958,616 | HiSeq 4000 |
| ATAC_Abd-B+Hth_n_105 | 26,325,518 | 19,788,572 | HiSeq 4000 |
| ATAC_Gcm_i_86 | 30,462,152 | 24,328,017 | HiSeq 4000 |
| ATAC_Gcm_i_87 | 18,888,376 | 15,277,774 | HiSeq 4000 |
| ATAC_Gcm_n_88 | 26,542,697 | 20,168,748 | HiSeq 4000 |
| ATAC_Gcm_n_89 | 12,643,646 | 9,371,579 | HiSeq 4000 |
| ATAC_Hth_i_58 | 31,491,896 | 22,619,681 | HiSeq 4000 |
| ATAC_Hth_i_59 | 27,549,490 | 20,662,463 | HiSeq 4000 |
| ATAC_Hth_n_60 | 31,034,936 | 23,887,936 | HiSeq 4000 |
| ATAC_Hth_n_61 | 34,868,300 | 27,248,785 | HiSeq 4000 |

**Table S4.** Hox group peak regions File available from https://doi.org/10.5281/zenodo.1491604. # Hox group peak regions for transient transfected samples, 15945 regions. # Location, data is dm6 release, start coordinates are 0-based # macs q-value 1e-2 columns are regions bound by both ChIP samples (low stringency) # macs q-value 1e-10 columns are regions bound by both ChIP samples high stringency) # Enhanced bound in Hox+Hth are regions more bound in the presence of Hth, edgeR fdr <= 0.01 and logFC > 1

**Table S5.** Cofactor-enhanced binding analysis of transient data in Hox group peak regions. # EdgeR results of ChIP-seq data locating enhanced bound regions in the presence of Hth using fdr <= 0.01 and logFC > 1. # Common bound are peak regions bound at similar levels (or do not have a significant fdr). # Hox enhanced bound (these may represent the noise in the genomic data) regions are fdr <= 0.01 but have a logFC < −1.

|  | Hox+Hth enhanced bound | Common bound | Hox enhanced bound |
| --- | --- | --- | --- |
| **Lab+Hth vs Lab** | **1248** | 2487 | 225 |
| **Pb+Hth vs Pb** | **463** | 5600 | 58 |
| **Dfd+Hth vs Dfd** | **1113** | 8264 | 239 |
| **Scr+Hth vs Scr** | **391** | 7769 | 94 |
| **Antp+Hth vs Antp** | **1140** | 1241 | 208 |
| **Ubx+Hth vs Ubx** | **428** | 3077 | 14 |
| **Abd-A+Hth vs Abd-A** | **282** | 5935 | 8 |
| **Abd-B+Hth vs Abd-B** | **351** | 9688 | 87 |

**Table S6.** Increased chromatin accessibility analysis. # EdgeR results of ATAC-seq data locating increased accessible regions in the 21,002 Hox group peaks - stable regions comparing induced (CuSO4) versus non-induced samples. # Increased accessiblity regions have fdr <= 0.01 and logFC > 1.5. # Decreased accessibility regions which have fdr <= 0.01 and logFC < −1.5 and may represent noise in the genomic data.

|  | increased accessibility | decreased accessibility |
| --- | --- | --- |
| Dfd induced vs Dfd non-induced | 430 | 122 |
| Ubx induced vs Ubx non-induced | 47 | 128 |
| Abd-B induced vs Abd-B non-induced | 832 | 103 |
| Hth induced vs Hth non-induced | 17 | 14 |

